# Demography and selection shape gene loss in freshwater sticklebacks - A pangenomic perspective

**DOI:** 10.64898/2026.09.08.750278

**Authors:** Hongbo Wang, Chaowei Zhang, Dandan Wang, Kerry Reid, Juha Merilä

## Abstract

The extent of within-species variation in genome assembly size, and the relative contributions of neutral *vs.* adaptive processes to it, remains poorly understood. We assembled the pangenome for the nine-spined stickleback (*Pungitius pungitius*) from whole-genome resequencing data obtained from range-wide sampling of 1,478 individuals across 47 populations. Our analysis recovered approximately 234.15 Mb of novel sequence and identified 11,151 protein-coding genes annotated in non-reference-represented regions, which accounted for 28.8% of all annotated pangenomic genes, corresponding to a 40.6% expansion relative to the 27,450 protein-coding genes annotated in the updated v8 reference genome. The identified presence/absence variation (PAV) genes exhibited a higher incidence of deleterious and non-synonymous mutations than core genes. Freshwater populations with smaller effective population sizes had lost more genes and had smaller assembly sizes relative to larger marine populations. Larger genome sizes of marine populations were primarily associated with higher transposable element content. Across gene categories, patterns of molecular variation were consistent with weaker purifying selection in freshwater populations than in marine populations, while recurrent freshwater-biased loss of a subset of PAV genes suggested that gene loss is not entirely random. Together, these results indicate that demographic history explains the overall pattern of gene loss, whereas negative natural selection may contribute to repeated loss of a subset of genes.

## Introduction

Pangenomes are large genomic datasets composed from hundreds or even thousands of individuals of a given species and are comprehensive representations of a species’ genome and diversity (Eizenga et al., 2020). Therefore, pangenomes offer a fuller picture of a species’ genetic diversity, including structural variation such as inversions, insertions, and deletions, and in particular the presence or absence of certain genes in different individuals or populations of the same species (Vernikos et al., 2015; Eizenga et al., 2020). The classic linear reference genomes for species are based on only one individual, and therefore cannot account for the wide breadth of single-nucleotide polymorphisms and structural variants within a species. Hence, access to a pangenome enables insights into genome evolution and local adaptation by enabling the capture of genomic variation hidden from the classic linear genomes (Tettelin et al., 2005).

Pangenomic resources have been developed for a limited number of species and are restricted by the availability of extensive high-quality resources. Vertebrate pangenomes are the most challenging to construct and have typically been constructed for humans (Miga & Wang, 2021; Wang et al., 2022; Liao et al., 2023; Wu Z et al., 2024) and other model and domestic species such as mice, pigs and chickens (Wang et al., 2021; Gong et al., 2023; Li et al., 2023; Helmy et al., 2026). However, the value of pangenomic tools is increasingly recognized as important for non-model species (Secomandi et al., 2025), and several pangenomes of wild vertebrates have been recently developed (Secomandi et al., 2023; Lan et al., 2024; Fang and Edwards, 2024; Quah et al., 2025). However, many of these have been based on a modest amount of data.

The nine-spined stickleback (*P. pungitius*), a small teleost fish, provides an ideal model for the development of pangenomic resources. This species is distributed circumpolarly across divergent marine and freshwater habitats in the northern hemisphere (Merilä, 2013), with the most basal lineage occurring in the western Atlantic (Guo et al., 2019; Fang et al., 2021). Nine-spined sticklebacks have repeatedly colonized freshwater habitats and become isolated from their marine ancestors. The freshwater populations show a wide range of effective population sizes (*N_e_*, Feng et al., 2025; Zhang et al., 2026) and correspondingly wide range of genetic diversity (Kivikoski et al., 2023a) and inbreeding levels (Kivikoski et al., 2023b; Chen et al., 2025), making them particularly interesting from evolutionary and pangenomic perspectives. In addition, the degree of genetic differentiation among nine-spined stickleback populations is much more pronounced than that of three-spined sticklebacks (Fang et al., 2021; Kemppainen et al., 2021), whereas the pool of standing genetic variation in nine-spined sticklebacks is much smaller and more fragmented than that of the three-spined stickleback (Fang et al., 2021, Kemppainen et al., 2021, Chen et al., 2025). In addition, two divergent lineages, western and eastern, inhabit Europe and hybridize in southern Scandinavia (Teacher et al., 2011; Feng et al., 2022; Yi et al., 2024). Hence, it is conceivable that genetic diversity at the pangenome level could be very high in the nine-spined sticklebacks.

Here, we describe the construction of the first linear pangenome for *P. pungitius* using a “map-to-pan” strategy (Hu et al., 2020) based on a large number of individuals (n = 1,478) from 47 populations of both marine and freshwater origins. Based on earlier observations on the degree of among population differentiation and within population genetic variability, we predicted that the pangenome would be substantially larger than the latest linear reference genome derived from a single freshwater individual (Wang et al., 2024). We further predicted that freshwater and marine populations would differ in gene content owing to a combination of demographic history, ecological divergence, and lineage specific gains and losses, and that populations with smaller *N_e_*s would show elevated gene loss. More specifically, we tested a framework in which genome-wide gene loss is shaped primarily by drift-driven fixation under reduced *N_e_*, whereas a narrower subset of recurrent freshwater-biased losses may reflect directional selection acting on particular genes. Under this framework, we expect a broad negative association between *N_e_*and gene loss across populations, but also a repeatable subset of losses shared across independent freshwater populations more often than expected by chance.

## Results

### The Pangenome and Comparison of Core and PAV Genes

The linear pangenome anchored to the v8 reference genome integrated *de novo* assemblies from 1,478 individuals representing 47 populations spanning marine, pond, lake, and river ecotypes (Supplementary Table 1 and Supplementary Table 1a). After filtering contaminated regions, we identified 234.15 Mb of novel contig sequences (n = 263,128) not present in the v8 reference genome. To improve the accuracy of identifying presence/absence variation (PAV) genes, we improved the quality of the reference genome annotation. The updated v8 gene annotation exhibited improved BUSCO completeness, with a score of 96.1% compared to the original 92.5% (Supplementary Table 2). Non-reference-represented (NRR) gene models showed a BUSCO score of 0.7%, indicating that our pipeline effectively reduced annotation errors attributable to allelic variation (Supplementary Table 2). The 11,151 NRR gene models refer to the full initial NRR annotation and correspond to a 40.6% expansion relative to the 27,450 protein-coding genes in the updated v8 reference annotation. After redundancy removal, quality filtering, and inclusion criteria for PAV genotyping, 8,726 NRR genes were retained in the final 36,174 gene PAV matrix; accordingly, the matrix used for downstream PAV analyses is not the simple sum of all initial v8 and NRR annotation models (Supplementary Tables 2 and 5).

The pangenome growth curve suggested saturation under the current sampling scheme, with the number of pangenes plateauing after approximately n = 682 sampled genomes and showing no marked increase beyond this point (Fig. 1a). In contrast, the size of the core gene set decreased with each added genotype, with no dramatic reduction after 392 accessions (Fig. 1a). The categorization of all 36,174 predicted genes by frequency revealed that 66.7% of the pangenome consisted of core genes (24,130 genes), with the remaining 33.3% (12,044 genes) comprising PAV genes that were further subdivided into dispensable (25.0%), softcore (5.6%), and private genes (2.7%) (Fig. 1b, Supplementary Table 3). The analysis of chromosomal distribution indicated that core genes were uniformly distributed across autosomes, accounting for approximately 90% of their gene content overall (range: 89.9% - 95.3% for LG1-LG21; Supplementary Table 4). In contrast, the X chromosome (LG12) had 86.8% of its gene content classified as core. We also found nine core genes in NRR regions that are absent from the v8 reference genome but present in all samples. These genes may correspond to regions collapsed in the v8 assembly. The pangenome helped recover and validate this information (Supplementary Table 5).

**Fig. 1.**
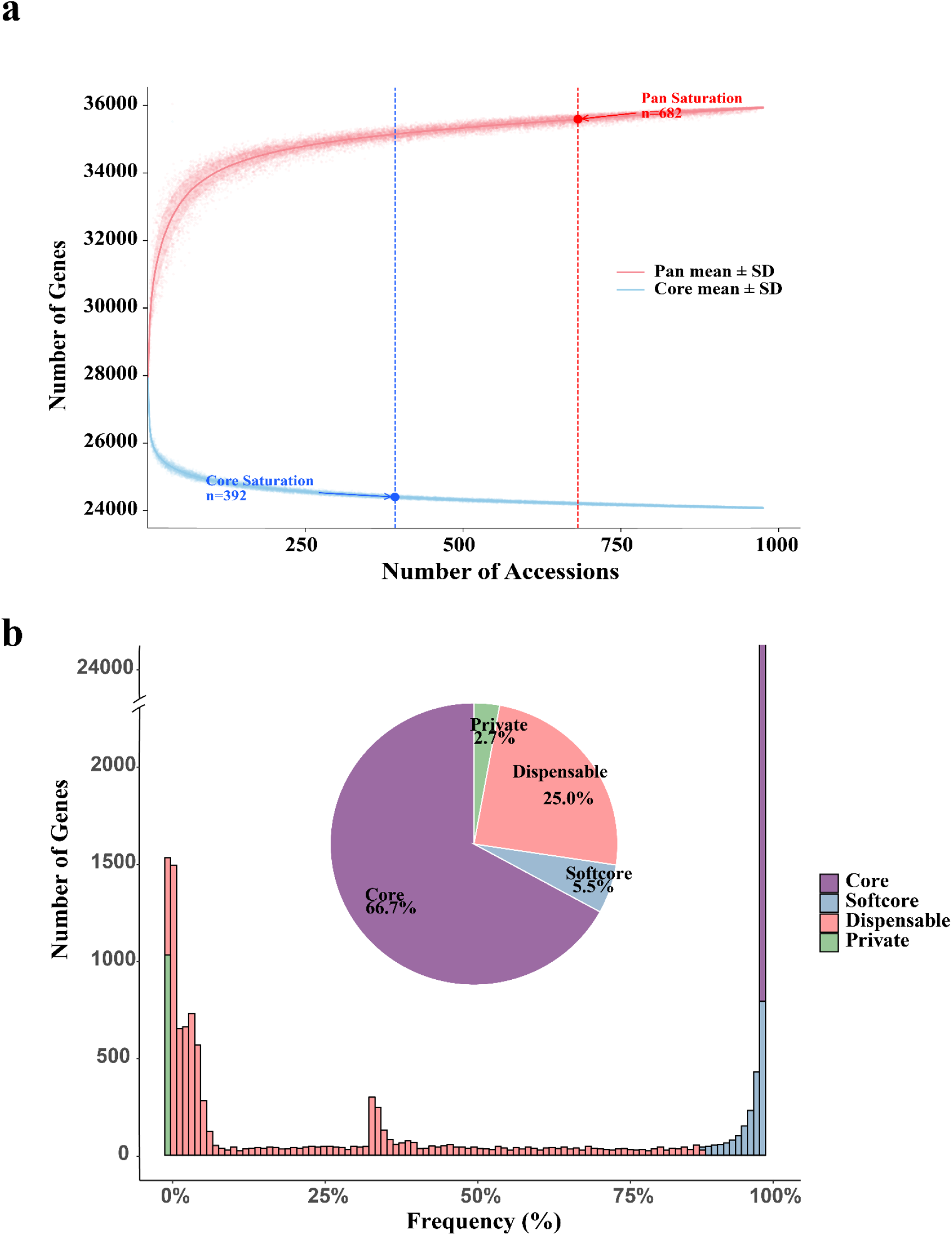
Pangenome of the nine-spined stickleback. (a) Modeling shows no marked increase in the number of genes after approximately n = 682 sampled genomes and no dramatic decrease in core gene numbers after n = 392. (b) Histogram and pie charts showing the frequency and number of core, softcore, dispensable, and private genes, respectively.

Gene Ontology (GO) enrichment analysis of 24,130 core genes indicated that basic physiological processes are conserved, with none of the biological processes significantly overrepresented (Supplementary Fig. 1). Among the 2,009 softcore genes, there was significant enrichment for functions related to RNA splicing (GO:0008380) and chromatin organization (GO:0006325) (q < 0.01) (Supplementary Fig. 2). Dispensable genes showed enrichment for several immune related functions, including interferon signaling (GO:0140888, q = 3.2 × 10^-5^) and MHC II antigen presentation (GO:0002399, q = 1.8 × 10^-4;^ Supplementary Fig. 3). Private genes also showed enrichment for pathways involved in B-cell regulation, specifically peripheral B-cell selection (GO:0002343, q = 4.7 × 10^-3^; Supplementary Fig. 4). Because immune related loci often show elevated copy-number variation, sequence divergence, and mapping complexity, we interpret these enrichments cautiously.

### Elevated Gene Loss in Small Freshwater Populations

Based on the set of 26,181 ancestral genes inferred from five ancestral populations, we detected significant differences in gene loss between freshwater and marine populations in the final ancestral gene loss subset of 809 individuals from 40 non-outgroup populations. The total gene number varied markedly among these populations, highlighting extensive gene PAV within the species (Supplementary Fig. 5). A simple comparison revealed that freshwater populations exhibited significantly greater gene loss than marine populations (t_551.01_ = 4.02, P = 6.7e-5; Fig. 2a), particularly in pond-dwelling populations (t_516.83_ = 8.00, P = 8.1e-15; Fig. 2b), among which the FIN-PYO population exhibited the lowest gene count of all (Supplementary Fig. 6).

**Fig. 2.**
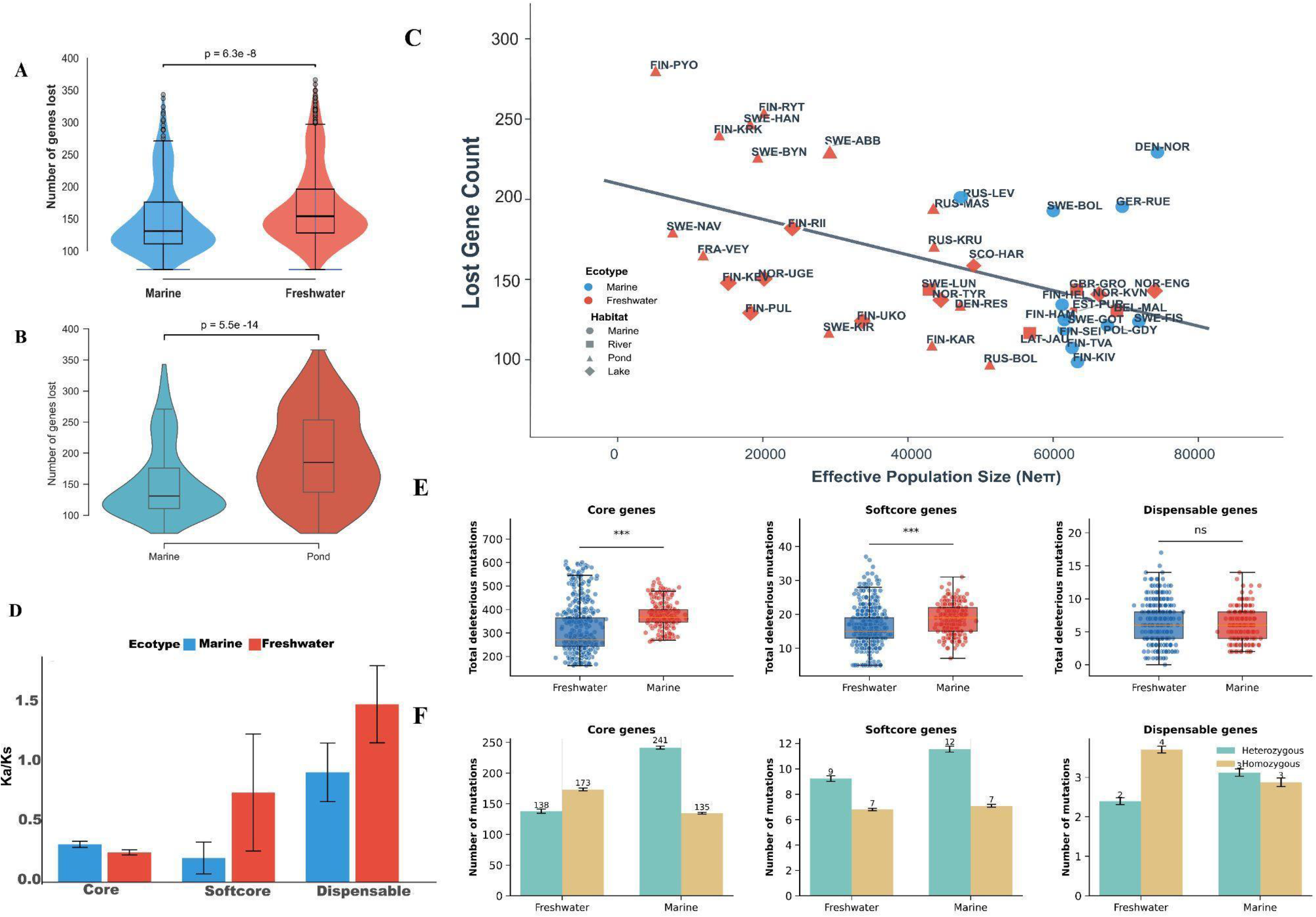
Gene loss and deleterious mutation load in freshwater and marine populations of nine-spined sticklebacks. (a) Average number of lost genes in freshwater and marine populations. (b) Average number of lost genes in pond and marine populations. (c) Relationship between effective population size (*N_e_*) and the number of lost genes. (d) Box plots showing nonsynonymous-to-synonymous ratio summaries in core genes and presence-absence variation (PAV) genes, grouped by ecotype. (e) Number of deleterious loss-of-function (LoF) mutations in core, softcore, and dispensable genes. (f) Number of deleterious homozygous and heterozygous LoF mutations in freshwater and marine populations.

To account for phylogenetic non-independence, we used a Bayesian phylogenetic Poisson mixed model with gene loss as the response variable, ecotype and *N_e_* as fixed effects, and phylogeny as a random effect. Gene loss decreased with increasing *N_e_* (*β* = -0.097, 95% confidence interval [CI]: -0.128 − -0.066, *p*MCMC < 0.001; incidence rate ratio (IRR) = 0.908, 95% CI: 0.879–0.937). After accounting for *N_e_*and shared evolutionary history, the ecotype effect changed direction so that the freshwater ecotype was associated with lower gene loss relative to the marine ecotype (*β* = -0.286, 95% CI: -0.342 − -0.229, *p*MCMC < 0.001; IRR = 0.751, 95% CI: 0.710–0.796). A sensitivity analysis including lineage as an additional fixed effect yielded similar estimates for habitat and *N_e_*, whereas lineage effects were weak and their 95% CI overlapped zero (Supplementary Table 7). This reversal indicates that the pronounced gene loss initially observed in freshwater populations compared to marine ones (Fig. 2a–b) is largely driven by the notably smaller *N_e_* inherent to freshwater habitats (Fig. 2c).

Population level sensitivity analyses yielded the same qualitative conclusion. Using the 40 populations with matched ancestral-gene-loss and *N_e_* information, mean gene loss per population remained negatively associated with log10(*N_e_* + 1), whereas the ecotype term was not significant after accounting for *N_e_*(Supplementary Table 7). Coverage based robustness analyses further showed that lower depth was associated with more inferred gene loss, but that the negative association between gene loss and *N_e_* remained significant after including either mean gene depth or the fraction of uncovered genes as additional technical covariates (Supplementary Table 7).

Together, these follow-up analyses support the same qualitative interpretation and are summarized in Supplementary Table 7. The ecotype effect was therefore model-dependent: it was weak or non-significant in the primary population level regressions after accounting for *N_e_*, although some rarefaction based summaries retained a directional difference. We therefore do not interpret the genome-wide pattern of gene loss as a simple freshwater-versus-marine contrast once demography is taken into account.

### Relaxed purifying selection in PAV genes and freshwater populations

We found that PAV regions exhibited elevated nonsynonymous-to-synonymous ratios (Fig. 2d). Specifically, freshwater populations showed significantly higher ratios in softcore and dispensable genes than marine populations, indicating reduced selective constraint on these genes (Fig. 2d). Marine and freshwater populations had similar ratios in core genes. There were insufficient data to estimate these ratios for private genes. The McDonald-Kreitman tests likewise supported weaker purifying selection in PAV genes than in core genes (Supplementary Fig. 7).

### PAV genes carry more deleterious variation

The nonsynonymous-to-synonymous ratio summary and the McDonald-Kreitman tests indicated that genes in PAV regions are under relaxed purifying selection. This relaxation is associated with an accumulation of loss-of-function (LoF) mutations in PAV regions (Fig. 2e). In unadjusted comparisons, considering total LoF variants (homozygous + heterozygous), marine populations showed significantly higher LoF burdens in core and softcore genes than freshwater populations (Wilcoxon rank-sum test, core: P = 2.26e-31; softcore: P = 2.82e-14), while dispensable genes had similar loads between ecotypes (P = 0.872; Fig. 2e). Because heterozygous LoF variants were less frequent in freshwater, we compared homozygous LoF counts: freshwater populations had more homozygous LoF genotypes in core genes than marine populations (mean: marine 143 vs. freshwater 184; P = 5.62e-41; Fig. 2e). This elevated homozygous LoF burden is likely driven by increased inbreeding and genetic drift in small freshwater populations, converting segregating deleterious variants into homozygous LoF burden. Softcore genes showed only a small difference across ecotypes (mean: marine 7.1 vs. freshwater 6.8; P = 9.85e-3), whereas dispensable genes carried slightly more homozygous LoF mutations in freshwater than in marine populations (3.7 in freshwater vs. 2.9 in marine; P = 1.7e-8; Fig. 2f). This also suggests that use of a single reference genome can bias estimates of mutation load.

### Non-random freshwater-biased gene loss

The repeatability analysis showed that recurrent freshwater-biased loss was more repeatable across populations than expected by chance. In the 39 population marine vs freshwater comparison, 86 genes were absent in at least 50% of individuals in at least half of freshwater populations but at most in one marine population, whereas the permutation null distribution had a mean of 1.61 such genes (empirical *P* = 0.001 based on 2,000 permutations; Supplementary Fig. 8). The pattern of loss was also highly parallel across independently derived freshwater populations (Supplementary Fig. 9). Supplementary Table 10 summarizes 44 genes from a broader recurrent loss screen used for descriptive follow-up; these genes do not necessarily meet the strict permutation-test criterion used for the 86-gene count. By contrast, the gene level *F_ST_* summary was not used as an independent inferential test, and none of the genes in the top 5% of the high-*F_ST_* distribution met our strongest repeatability criterion. For the subset of recurrent-loss candidates that could be traced directly in the final ancestral-loss framework, exon-level coverage summaries showed that freshwater LOST calls corresponded to near-complete exon-wide coverage collapse, whereas marine PRESENT calls at the same loci retained high exon coverage (Supplementary Table 8). Together, these results suggest that gene loss in freshwater populations is not entirely random and that a subset of recurrent losses may reflect shared ecological selective pressures, although demographic history and technical complexity remain important caveats to this interpretation.

### Assembly-span variation across pond and marine populations

To examine possible differences in genome size between pond and marine ecotypes, we *de novo* assembled genomes of 434 individuals from four populations of each ecotype. Analysis of the resulting assembly sizes disclosed a clear pattern: marine individuals consistently yielded larger assembly-based genome spans than pond individuals (Table 2, Fig. 3a). Specifically, the mean genome assembly size for marine populations ranged from 462.4 Mb to 474.0 Mb, whereas those of pond populations ranged from 441.2 Mb to 448.6 Mb. This direction was independently supported by a fully phased T2T comparison between a marine TVA assembly and the freshwater PYO reference, after removing chrY, the TVA assembly spanned 464.65 Mb excluding mitochondrial sequence, matching the range estimated from the independent short-read marine assemblies (462.4-474.0 Mb). Under the more conservative homologous non-sex-chromosome comparison, anchored sequences totaled 429,327,121 bp in TVA but 406,173,618 bp in PYO, and the PYO total reached only 409,127,584 bp even when unanchored contigs were included (Zhang et al., 2026). The quality of these assemblies was high in terms of completeness. Initial contigs generated from short-read data achieved BUSCO completeness scores of 82.6-88.1% across populations. Duplication rates were low (0.56-0.75%), suggesting that excessive duplication is unlikely to be the sole explanation for the larger genome assembly sizes observed in marine populations. These contigs were subsequently scaffolded against the v8 reference genome to produce chromosome-level assemblies, which increased the BUSCO scores to 98.15-98.35% (mean 98.2%, Fig. 3b), while duplication remained low (0.57-0.76%; Table 2, Fig. 3c). These findings indicate that the assemblies are of comparable overall quality and suitable for comparative genome-assembly-size analyses. To investigate factors underlying among-population differences in genome assembly size, we fitted linear mixed-effects models (LMMs). First, repeat content emerged as the strongest predictor of assembly size in a mixed model with population as a random intercept (Model 1: Assembly_Size ∼ Repeat_Content_scaled + (1|Population); P < 1e-16), consistent with the expectation that repeats dominate genome-scale variation.

**Fig. 3.**
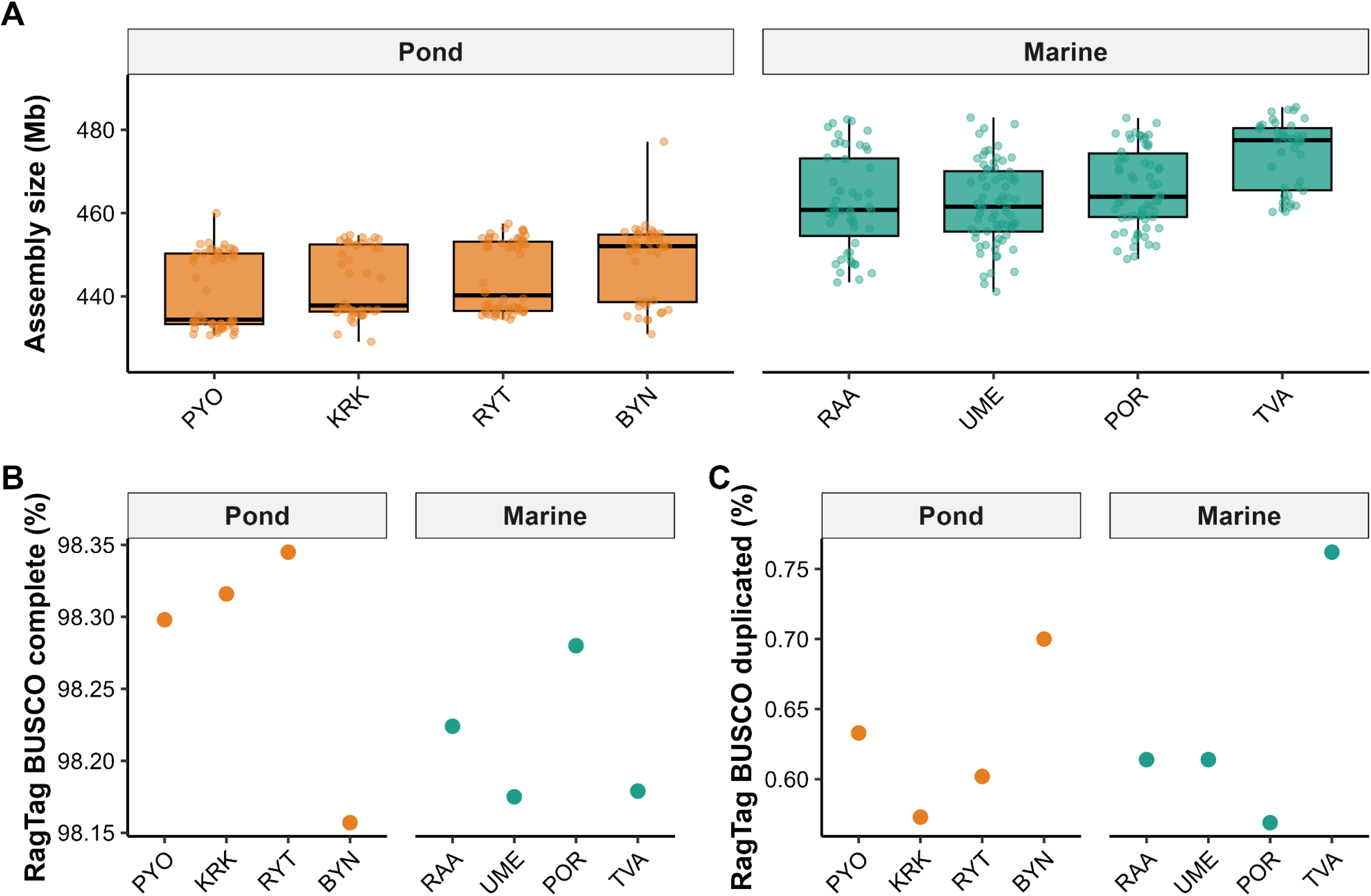
Genome-assembly-size distribution and BUSCO scores replicate populations of pond and marine nine-spined sticklebacks. (a) Genome assembly size for individuals from each population. (b) Mean percentage of complete BUSCOs in RagTag assemblies for each population. (c) Mean percentage of duplicated BUSCOs in RagTag assemblies for each population.

**Table 1.** A summary of the results of the MCMCglmm model for gene loss. IRR = incidence rate ratio.

| Fixed Effects (Predictors) | Posterior Mean ($\beta$) (95% CI) | IRR(95% CI) | $P_{\text{MCMC}}$ |
| --- | --- | --- | --- |
| Intercept | 4.931 [4.463, 5.392] | 138.485 [86.753, 219.575] | < 0.001<br>*** |
| *Ne* (log scaled) | -0.097 [-0.128, -0.066] | 0.908 [0.879, 0.937] | < 0.001<br>*** |
| Ecotype | -0.286 [-0.342, -0.229] | 0.751 [0.710, 0.796] | < 0.001<br>*** |

**Table 2.** A summary of genome assembly size and BUSCO completeness in eight nine-spined stickleback populations.

| Populat<br>ion | Ecoty<br>pe | Sam<br>ple<br>size<br>(n) | Mea<br>n<br>geno<br>me<br>size<br>(Mb) | SD<br>geno<br>me<br>size<br>(Mb) | 95%<br>CI of<br>geno<br>me<br>size<br>lowe<br>r<br>(Mb) | 95%<br>CI of<br>geno<br>me<br>size<br>uppe<br>r<br>(Mb) | Assem<br>bly<br>BUSC<br>O<br>compl<br>ete<br>(%) | Assem<br>bly<br>BUSC<br>O<br>duplica<br>ted (%) | RagT<br>ag<br>BUSC<br>O<br>compl<br>ete<br>(%) | RagTa<br>g<br>BUSC<br>O<br>duplica<br>ted (%) |
| --- | --- | --- | --- | --- | --- | --- | --- | --- | --- | --- |
| BYN | Pond | 44 | 448.58 | 9.41 | 445.72 | 451.44 | 85.81 | 0.66 | 98.15 | 0.70 |
| KRK | Pond | 44 | 443.19 | 8.35 | 440.66 | 445.73 | 86.66 | 0.61 | 98.32 | 0.57 |
| PYO | Pond | 55 | 441.16 | 9.04 | 438.72 | 443.61 | 88.14 | 0.64 | 98.30 | 0.63 |
| RYT | Pond | 60 | 444.73 | 8.54 | 442.52 | 446.94 | 85.03 | 0.56 | 98.35 | 0.60 |
| POR | Marine | 66 | 466.83 | 12.23 | 463.82 | 469.84 | 83.55 | 0.62 | 98.28 | 0.57 |
| RAA | Marine | 51 | 462.40 | 11.54 | 459.15 | 465.64 | 83.05 | 0.65 | 98.22 | 0.61 |
| TVA | Marine | 42 | 474.03 | 8.27 | 471.45 | 476.61 | 83.97 | 0.75 | 98.18 | 0.76 |
| UME | Marine | 72 | 462.67 | 11.06 | 460.07 | 465.27 | 82.56 | 0.64 | 98.18 | 0.61 |

CI, confidence interval.

Next, to test whether gene loss is associated with assembly contraction, we fitted a model excluding repeat content to avoid collinearity. The number of lost PAV genes was strongly and negatively associated with assembly size (Model 2: Assembly_Size ∼ Gene_Loss_scaled + (1|Population); P < 1e-9), indicating that populations with greater gene loss also tended to yield smaller short-read genome assemblies.

A joint model including repeat content and gene loss showed severe collinearity between these predictors (VIF > 15). We therefore do not use that joint model as a basis for primary inference or variance partitioning and instead interpret repeat content and gene loss as correlated predictors of genome-assembly-size variation.

## Discussion

There are many theories of genome size evolution, some of which emphasize the role of neutral processes, and others which emphasize the role of natural selection (Blommaert 2020; Marino et al., 2024). The most salient finding of our intraspecific study is that both demographic history and selection contribute to genome content and genome size variation in nine-spined sticklebacks. We found strong associations among ecotype, effective population size (*N_e_*), gene loss, repeat content, and genome assembly size, but the observed associations do not support a single genome-wide mechanism. Instead, our results suggest that the overall excess of gene loss in many freshwater populations is primarily associated with reduced *N_e_*, whereas repeated loss of a subset of genes across freshwater populations is consistent with non-random recurrent change that may involve selection. In the following, we discuss these findings and how they contribute to our understanding of genome content variation and genome size evolution.

A key finding of our intraspecific study is that genome-assembly-size variation does not conform neatly to the classic expectation (Lynch and Conery 2003) where reduction in *N_e_*leads to weakened purifying selection, transposable element (TE) accumulation, and subsequent genome expansion. In contrast, we found that large-*N_e_* marine populations had much higher TE burdens than smaller freshwater populations. This was unexpected because small freshwater populations subject to weaker purifying selection might be expected to accumulate more TEs and therefore have larger assemblies than large marine populations. One possible explanation for this counterintuitive observation lies in the demographic history of freshwater populations. If freshwater populations were established by a small number of founders that happened to carry fewer copies of TE families present in the ancestral population, the newly founded populations would start with a lower TE load simply due to random sampling of the initial gene pool. Also, since the freshwater populations were established less than 10,000 years ago, there might not have been sufficient time for TE families to proliferate. Furthermore, the strong geographic isolation of freshwater populations limits gene flow and reduces their ability to acquire new active TE families through exchange among populations, unlike marine populations. In addition, if most TEs already present in freshwater populations are inactive, these genomes are less likely to produce new insertions, leading to a much lower TE load than in marine populations. Yet another possible explanation for low TE burden in freshwater populations is that repeat landscapes may turn over rapidly over short evolutionary timescales owing to founder effects, stochastic losses of TEs, and differences in repeat turnover among populations.While we cannot currently differentiate in between these alternative explanations, it is clear that the lower TE burden contributes to the smaller genome assemblies in freshwater populations.

The genome streamlining hypothesis (Blommaert 2020) suggests that limited resources, or selection for metabolic efficiency or improved cell division, may favor smaller genome sizes (Blommaert 2020). However, our results indicate that purifying selection was not stronger in the small freshwater populations than in the large marine populations. Consequently, our findings do not support the idea that the overall reduction in genome assembly size is a simple adaptive response driven by more efficient genome-wide removal of deleterious variants. Instead, this reduction is better explained by the populations’ demographic histories and natural fluctuations in repetitive region abundance. At the same time, our analyses suggest that the independent and repeated loss of a subset of specific genes may be non-random and could reflect shared ecological selective pressures in freshwater environments. Ultimately, while the data are consistent with non-random loss of particular genes, they do not fully explain contraction of the entire genome. Whatever the cause of smaller assemblies in small-*N_e_*freshwater populations, our results align with the conclusion of the recent comparative study finding no evidence that increased drift necessarily leads to the accumulation of TEs across animals (Marino et al., 2024). However, our results are also consistent with the recent findings of Edwards et al. (2025), who found that over relatively short evolutionary timescales, genome size dynamics can be driven by rapid turnover of repeat landscapes: lineages with the lowest *N*_e_s had the smallest genomes, challenging the classic view (Lynch and Conery 2003; Lynch 2007).

Genome contraction in freshwater populations involved not only a reduction in TE content, but also a decrease in gene number. The elevated loss among presence-absence variation genes suggests that this process contributes to genome content reduction. We found that although freshwater populations exhibited substantial genomic variation, some aspects of gene loss were non-random. In particular, recurrent freshwater-biased losses were more repeatable across populations than expected under random ecotype assignment, and some of these losses reoccurred across multiple independent freshwater populations that do not trace back to a single ancestral coalescent event. For the recurrent-loss candidates represented in the ancestral-loss framework, exon-level coverage patterns were also consistent with true loss rather than patchy borderline undercoverage. This recurrent pattern is consistent with the possibility that some gene losses are tolerated and repeatedly fixed in freshwater environments, although the evidence does not justify attributing the full genome-wide pattern of gene loss to selection alone. Similar findings have been reported for the common bean *(Phaseolus vulgaris*), where pangenome contraction occurred during range expansion and domestication and was interpreted in that system as involving adaptive gene loss (Cortinovis et al., 2024). This comparison suggests that selection can contribute to gene loss in some contexts, but it does not by itself demonstrate selection in nine-spined sticklebacks. Hence, we propose that among-population variation in genome content in nine-spined sticklebacks can be understood in light of at least two processes. First, TEs and other repetitive sequences are the primary determinants of overall genome assembly size. Second, population-specific gene loss, shaped by both demographic history and selection on a subset of genes, contributed to reduced gene content in many freshwater populations. When sticklebacks colonized freshwater habitats, they likely experienced severe bottlenecks that reduced *N_e_*, creating conditions that favor strong genetic drift (Feng et al., 2022). Although drift can cause random gene loss, our data suggest that not all losses are distributed uniformly across the genome.

At the same time, our follow-up sensitivity analyses indicate that this genome-wide pattern should not be reduced to a single freshwater-versus-marine adaptive contrast. The direction of the population-size effect was stable across alternative *N_e_* estimators, count-model formulations, and rarefaction analyses, whereas local marine-freshwater contrasts were significantly heterogeneous across geographic clusters. We therefore interpret the recurrent freshwater-biased losses as a narrower pattern superimposed on a broader demographic background, rather than as evidence that freshwater habitat alone drives the total excess of gene loss across the genome.

The western (WL) and eastern European lineages (EL) of nine-spined sticklebacks have different and independent evolutionary histories (Feng et al., 2022), including different sex chromosomes and sex-determination systems (Yi et al., 2024). Hence, our expectation was that there would also be marked differences in gene content between the two lineages. However, comparison of gene numbers between the two lineages revealed fairly similar gene numbers: WL had 31,202 genes whereas EL had 33,714 genes. No fully fixed diagnostic PAVs were identified between the lineages, indicating limited fixed gene content divergence. More broadly, many PAVs appear to be restricted to one lineage without being fixed within that lineage, which is consistent with recent, lineage-specific duplication and loss events rather than simple retention of deep ancestral variation. By contrast, the absence of fixed PAVs between the lineages suggests that the strongest repeated gene-loss patterns discussed above are not solely a lineage-specific phenomenon.

While the application of pangenomics to natural animal populations is rapidly expanding, most vertebrate studies remain constrained by limited sample sizes, which makes our study one of the most comprehensively sampled pangenomic studies of any non-model species (Supplementary Table 9). By operating at a scale substantially larger than typical wild vertebrate pangenome studies, we were able to make robust inferences regarding population-level presence-absence variation. We found that 33% of the *P. pungitius* gene pool consisted of PAVs. When compared to existing literature, this level of structural plasticity is exceptionally high. The observed frequency of PAV genes is comparable to plants (e.g., ∼38% in cultivated rice; Wang et al., 2018) but largely unseen in vertebrates, whose genomes are typically considered stable (Brůna et al., 2026). Our findings therefore suggest that a large component of dispensable genes is not exclusive to plants or microbes. More broadly, extensive PAVs may characterize vertebrate populations with complex demographic histories, fragmentation, and repeated environmental transitions.

The results also illustrate limitations of traditional reference genome assemblies. The most recent nine-spined stickleback genome assembly (v8; Wang et al., 2024) is the most contiguous and complete available before this study. However, the individual used in its assembly came from the most homozygous and smallest *N_e_* population among the 47 *P. pungitius* populations studied (FIN-PYO; Shikano et al., 2010; Chen et al., 2025) greatly restricting its coverage of ancestral genetic variation. This was one of the motivations for adopting a map-to-pan strategy rather than relying on the original reference assembly alone. For comparison, the v8 reference genome (FIN-PYO population) showed 99.6% complete BUSCOs and 3.4% duplication in the assembly-level assessment, whereas our independent FIN-PYO population assembly (estimated size 441.16 Mb) achieved 98.3% completeness and 0.63% duplication in the corresponding assembly-level assessment.

This within-population comparison supports the view that repeat collapse in the reference assembly, rather than population identity alone, contributes to the reduced span of the published PYO reference. The same conclusion is also supported by an independent fully phased T2T marine TVA assembly, which exceeds the freshwater PYO reference in total span (Zhang et al., 2026). In addition, as revealed by our pangenome assembly, aggressive repeat collapsing during the v8 assembly produced artificial losses of repetitive DNA, leading to a distorted view of the *P. pungitius* genome. These shortcomings not only hide potentially interesting variation, but can also lead to biased inferences, as demonstrated by the apparent enrichment of loss-of-function mutations in PAV genes when analyses rely on a single reference genome.

The newly assembled linear pangenome leaves room for several questions and methodological limitations to be addressed in future studies. First, linear pangenomes have lower accuracy than graph pangenomes for detecting complex features of the data. However, the linear approach was chosen due to the considerable cost of performing long-read and Hi-C sequencing on hundreds of individuals. Second, limited by short-read sequencing technologies, accurate cross-population comparison in extremely complex genomic regions such as segmental duplications (SDs) and low-complexity regions continues to be an issue. This will also reduce the accuracy of site frequency spectrum (SFS) analyses. Moreover, the immune-related enrichments reported above point to complex loci such as the MHC as promising targets for future work; haplotype-resolved genomes will be needed to detail the structural diversity of such highly polymorphic regions. Third, the timing of TE insertions needs to be studied further. Fine-scale annotation and eventual comparison of TE insertions across different families of large-*Ne* marine populations and small *N_e_* freshwater populations can uncover possible differences in their TE evolutionary dynamics. On a methodological front, pedigree-based long-read sequencing will enhance *de novo* TE detection. This will facilitate comparative studies to reveal whether small populations experience more frequent *de novo* TE insertions, even though their total TE accumulation is smaller. Finally, while our findings suggest that a subset of PAV genes may be associated with shared ecological or selective pressures in freshwater habitats, the phenotypic and ecological consequences of these gene losses remain unknown. Integrating pangenomic data with gene-expression profiling, functional assays, and environmental measurements should help clarify the biological drivers consequences of recurrent PAV loss across freshwater populations.

In conclusion, the *P. pungitius* pangenome shows that genome content and short-read genome-assembly-size variation are shaped by a complex interplay between demography and selection. Our results indicate that freshwater colonization in this species involved not only allelic divergence but also substantial remodeling of the gene repertoire, with overall gene-loss patterns strongly associated with demographic history and a subset of recurrent losses suggesting non-random change across freshwater populations. These findings underscore the value of pangenomic approaches in evolutionary biology, particularly for species with high intraspecific diversity and complex demographic and ecological histories.

## Materials and Methods

### Re-sequenced genomes and populations

In this study, we used 1,478 re-sequenced *P. pungitius* genomes from previously published datasets including 887 wild samples from 45 populations sequenced to an average depth of approximately 10x using standard Illumina short-read whole-genome resequencing (paired-end reads on HiSeq 2500/4000; Feng et al., 2022; Kivikoski et al., 2023a), 157 additional individuals from the FIN-HEL population sequenced to approximately 5x depth (Kivikoski et al., 2021), and 434 individuals from eight populations representing pedigreed individuals sequenced using PCR-free DNBseq to an average read depth of 80x (Zhang et al., 2023; Zhang et al., 2026). As some populations were represented in more than one dataset, the total number of unique populations included in this study was 47, with sequencing depth ranging from approximately 5× to 10× in the population-level datasets and averaging 80× in the family-based dataset. The sampled populations included both freshwater and marine populations, and information on sampling site coordinates and population specific sample sizes is given in Supplementary Table 1 and the analytical use of each sample set is summarized in Supplementary Table 1a.

### Pangenome construction

The workflow for the “map-to-pan” method for constructing the linear pangenome is provided in the Supplementary Material. We first employed fastp v0.23.4 (Chen et al., 2018) to remove adapters and low-quality bases from the raw sequencing data. After quality control, all 1,478 samples were de novo assembled. The 1,044 individuals from the population-level datasets (887 wild individuals from 45 populations plus 157 FIN-HEL individuals) were assembled de novo using SOAPdenovo2 (Luo et al., 2012). Any gaps remaining in the scaffolds generated by SOAPdenovo2 (Luo et al., 2012) were closed using GapCloser (Luo et al., 2012) to improve contiguity. For the data from eight populations with high sequence depth (Zhang et al., 2023; Zhang et al., 2026), all 434 samples were suitable for draft genome-size estimation and were assembled separately using MEGAHIT (Li et al., 2015) with multiple k-mers (21, 29, 39, 59, 79, 99, 119, and 141). Each assembly was then scaffolded to the chromosome level using RagTag v2.1.0 (Alonge et al., 2022) by aligning it to the v8 reference genome, which was derived from the highly homozygous freshwater population FIN-PYO (Wang et al., 2024). The resulting contig-level assemblies were aligned to the v8 reference genome, and non v8 reference contigs that were unaligned and longer than 500 bp were extracted based on QUAST (Gurevich et al., 2013) alignment results. Owing to the prolonged runtime of CD-HIT-EST v4.8.1 (Li et al., 2006), we initially employed MMseqs2 (Steinegger and Söding, 2017) to prefilter the sequences. Next, we applied CD-HIT-EST v4.8.1 (Li et al., 2006) with a default global sequence-identity threshold of 90%, calculated as the number of aligned bases divided by the full length of the shorter sequence to eliminate redundant sequences that could not be mapped to the v8 reference genome. The filtered sequences were then aligned against the NCBI Nucleotide database using BLAST (Altschul et al., 1990). Only contigs identified as animal-derived were retained and incorporated into the pangenome as non-reference representative sequences. Finally, Kraken2 (Wood et al., 2019) and Minimap2 (Li et al., 2018) were used to filter out mitochondrial genomes from the reference dataset, and only contigs of at least 500 bp in length were retained. Thus, the downstream analyses were based on a v8-anchored linear pangenome that combined the original reference component with non-reference representative sequences recovered across 1,478 individuals, rather than on the original FIN-PYO reference genome alone.

### Pangenome annotation and evaluation

To annotate the pangenome genes, we first soft-masked transposable elements (TEs) in the linear pangenome. RepeatModeler (Flynn et al., 2020) was used for de novo discovery of TE families, and RepeatMasker (Tarailo-Graovac et al., 2009) identified repetitive elements by mapping them to the RepBase database (Bao et al., 2015) using a homology-based method. We performed separate annotations for the reference genome and the non-reference representative region (NRR). To update the reference genome annotation (version 8; Wang et al., 2024), we utilized the BRAKER v2.1.6 pipeline (Brůna et al., 2021) with protein sequences from SwissProt database (zebrafish: 3,683; Japanese killifish: 93; three-spined stickleback: 4; total = 3,780 genes) and transcript data from Wang et al. (2020) from six populations (n = 24, with two males and two females per population). The RNA-seq data from brain and liver were mapped to the v8 reference genome using HISAT2 (Kim et al., 2019). The initial BRAKER predictions were then refined with the PASA pipeline (Haas et al., 2003), which adjusted exon boundaries, and added untranslated regions. Additionally, we used SynGAP (Wu F et al., 2024) to polish and refine the annotations with annotation data from Danio rerio (GCA_000002035.4) and Gasterosteus aculeatus (GCA_016920845.1) in the Ensembl database.

For NRR annotation, we used AUGUSTUS (Stanke et al., 2004) to carry out ab initio annotation of novel regions with a gene model trained from the output of BRAKER in the reference-annotation step described above. We also generated homology-based annotations for the same protein-coding genes sourced from the SwissProt database using miniprot (Li, 2023). We then used the HISAT2 alignments of the same transcript dataset to assemble transcripts with StringTie (Pertea et al., 2015) and combined *ab initio* gene predictions with protein and transcript alignments into weighted consensus gene structures. In EVidenceModeler (EVM; Haas et al., 2008), the weights were set to 1 for *ab initio* gene predictions, two for miniprot protein alignments, and eight for transcript alignments. We then compared the EVM predictions with the StringTie assemblies. For any gene annotation from EVM with a coding sequence (CDS) coverage below 30%, we performed a Pfam search using the HMM database via hmmscan (Eddy, 2011), with an E-value threshold below 1 × 10^-6 and a minimum coverage of 30%. Gene models with no significant matches in the Pfam search were discarded. The Pfam search was carried out using hmmscan with the Pfam-A HMM database and the filtering script available from https://github.com/Zhanmengtao/bin/blob/master/evm_genes_filtering.pl. To evaluate the new annotation quality of the v8 reference assembly, we used BUSCO v6.0.0 (Manni et al., 2021) to assess gene-annotation completeness against the actinopterygii_odb10 single-copy gene database. We also evaluated the non-reference representative sequence annotation for redundancy and allelic gene annotations.

### PAV calling and analyses

After assembling and annotating the linear pangenome, 967 individuals with sufficient depth for reliable PAV genotyping were aligned using BWA-MEM2 (Vasimuddin et al., 2019) with default parameters. These 967 individuals comprised the 887 wild samples from 45 populations and 80 F0 individuals from the high-depth pedigree dataset. F1 and F2 individuals were excluded because they are not independent population samples. This complete set of 967 samples was used to model the dynamics of the pangenome. The presence or absence of genes was determined using SGSGeneLoss software (Golicz et al., 2015). For the high-depth sequencing dataset from eight populations, F0 samples (80 individuals; Zhang et al., 2023; Zhang et al., 2026) were retained and stricter parameters were applied (minimum coverage = 10, loss cutoff = 0.2). For the dataset comprising 887 samples from 45 populations (Feng et al., 2022), a gene was classified as absent if less than 20% of its exon regions were covered by at least two sequencing reads (parameters: minimum coverage = 2, loss cutoff = 0.2). The same parameter combination has been used in several SGSGeneLoss based PAV studies in plants (Gao et al., 2019; Li et al., 2021; Barchi et al., 2021), but because some of those studies used higher sequencing depth than the present wild dataset, we did not rely on literature precedent alone. Instead, these parameter settings were calibrated primarily from empirical comparisons showing that PAV calls were consistent within each sequencing depth class. To minimize bias arising from pedigree structure and uneven sequencing depth, all subsequent population-level PAV and coverage analyses were restricted to 887 wild, unrelated individuals sequenced at broadly comparable depth. PAV results for these 887 wild samples were then summarized across the two divergent lineages and ecotypes.

The sizes of the pangenome and core genome were estimated using a custom script by randomly selecting 2,000 different orders for the 967 samples (https://github.com/LeoHongboWANG/sticklebacks_PAV). These models were fitted using nonlinear least-squares regression in R with the nlsLM function. The expansion of the pangenome was modeled using a power-law function:

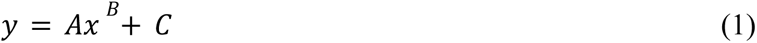

The reduction in core genome size was modeled using an exponential decay function:

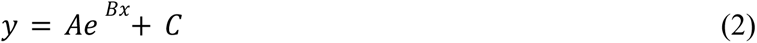

Based on the PAV results, genes were classified into four categories: (1) “Core genes” present in all samples (100%); (2) “Softcore genes” present in 90–99% of samples; (3) “Dispensable genes” present in fewer than 90% but in at least two samples; and (4) “Private genes” present in only a single sample.

To infer the ancestral gene set, we used five populations from Canada, Russia, the USA, and Japan (CAN-FLO, CAN-TEM, JAP-BIW, RUS-LEN, and USA-HLA) as outgroups, which included representatives from both the Eastern (EL) and Western Lineage (WL) ancestral datasets (Feng et al., 2022). Genes present in more than 90% of outgroup individuals were designated as the ancestral gene set. We then quantified gene losses for EL and WL samples relative to the ancestral gene set, excluding instances in which ancestral genes were absent.For binary ecotype analyses, pond, lake, and river populations were grouped as freshwater, whereas marine populations were treated as the marine category. The final ancestral-gene-loss dataset therefore comprised 809 wild, unrelated individuals, from 40 non-outgroup populations.

### Gene Ontology (GO) enrichment analysis

GO annotation of the pangenome was performed using genes that were functionally annotated with eggNOG v5 (Huerta-Cepas et al., 2019) with an E-value cutoff of 1 × 10^-5. GO enrichment analysis was conducted with the R package clusterProfiler v3.14.3 (Yu et al., 2012), using all coding genes as the background.

### SNP calling

Short-read sequencing data from the 887 wild individuals were aligned to the linear pangenome using BWA-MEM2 (Vasimuddin et al., 2019). BAM files generated from the presence/absence calling step were sorted using SAMtools (v1.18; Li et al., 2009), and PCR duplicates were marked using Sambamba (Tarasov et al., 2015). Variant calling was carried out using GATK HaplotypeCaller (Van der Auwera & O’Connor, 2020) to generate individual genomic variant call format (gVCF) files. These gVCF files were jointly genotyped using GLnexus (Yun et al., 2021). We performed genotyping on gene regions, applying the filtering strategy described by Feng et al. (2022) (https://github.com/XueyunF/nsp_phylogeo). Briefly, intergenic variants were excluded, and biallelic SNPs were identified using BCFtools v1.7 (Danecek et al., 2021). Sites with low average coverage (<8×), genotype quality (<20), or overall quality score (<30) were filtered out using VCFtools v0.1.17 (Danecek et al., 2011).

### Synonymous and nonsynonymous variation and detection of deleterious variation

To identify deleterious variants, we adopted the approach of Chen et al. (2025) to polarize SNPs using the ancestral EL and WL population RUS-LEN (Feng et al., 2022), with a customized script (https://github.com/LeoHongboWANG/sticklebacks_PAV). At each site, the reference allele (REF) was considered ancestral when at least 80% of the outgroup samples were homozygous reference (0/0); if at least 80% were homozygous for the alternative allele (1/1), we inverted ALT to REF. Using the adjusted reference alleles, we generated an updated reference genome using GATK’s FastaAlternateReferenceMaker module, representing the ancestral state of the *P. pungitius* genome. We used degenotate v1.3 to summarize synonymous and nonsynonymous coding-site classes, which were then used for the gene-level McDonald-Kreitman (MK) summaries and estimating the nonsynonymous-to-synonymous ratios. This modified genome also served as the basis for subsequent variant annotation performed with SnpEff (Cingolani et al., 2012), ensuring consistency in allele labeling across analyses.

### Statistical model comparisons

To compare gene loss between the EL and WL lineages, as well as between ecotypes, we employed Bayesian phylogenetic Poisson mixed models using MCMCglmm (Hadfield, 2010). Gene loss was modeled as the response variable. In the final main model, fixed effects included scaled log10(*Ne* + 1) and ecotype, with the Marine set as the reference level. To account for phylogenetic non-independence among samples, phylogeny was included as a random effect using the VCF2Dis tree (Xu et al., 2025). Lineage was not retained as a fixed effect in the main model because lineage-level structure is largely captured by the phylogenetic random effect, including the major divergence between the EL and WL lineages; instead, lineage was included only in a sensitivity analysis, which tested whether the main results were robust to the inclusion of lineage as an additional fixed effect. Populations identified as hybrids between the EL and WL lineages had been excluded during data preprocessing, based on Feng et al. (2022) and Yi et al. (2024). Models were run with four independent MCMC chains (nitt = 1,300,000; burn-in = 300,000; thinning interval = 500), and posterior summaries were checked for consistency across chains before inference. Additional population level and coverage based sensitivity analyses are summarized in the Supplementary Table 7.

To evaluate whether recurrent freshwater biased gene loss exceeded random expectation, we performed a population level repeatability analysis using the revised marine vs freshwater comparison (27 freshwater and 12 marine populations). For each gene, we counted the number of populations in which at least 50% of individuals lacked the gene. We defined strongly repeatable freshwater-biased loss as genes absent in at least half of the freshwater populations but in at most one marine population, and compared the observed number of such genes with a null distribution generated by 2,000 permutations of ecotype labels across populations. In parallel, we calculated gene-level FST between marine and freshwater populations from the PAV matrix. Presence and absence were encoded as pseudo-genotypes (1/1 and 0/0, respectively), the resulting matrix was converted to VCF format, and FST was estimated using VCFtools v0.1.17 (Danecek et al., 2011). Because differentiation (FST) and freshwater-biased loss are not fully independent, we treat the gene-level FST comparison as descriptive rather than as a stand-alone test of selection.

## Acknowledgments

We thank all people who over the years have provided the samples used in this study, and in particular, Pär Byström, Xueyun Feng, Antoine Fraimout and Ulrika Candolin. We acknowledge the Finnish IT Centre for Scientific Computing (CSC) for access to high-performance computational resources. Our research was funded by the Research Grants Council of Hong Kong (General Research Fund: 17126824 to KR & JM).

## Code availability

The codes used in this paper are available on GitHub (https://github.com/LeoHongboWANG/sticklebacks_PAV).

## Data availability

The pangenome assembly and annotation are available at Zenodo (10.5281/zenodo.18591803).

