## Supplemental Information for "Demography and selection shape gene loss in freshwater sticklebacks - A pangenomic perspective"

**Supplementary information**

**Supplementary Fig. 1.** The GO enrichment of core genes

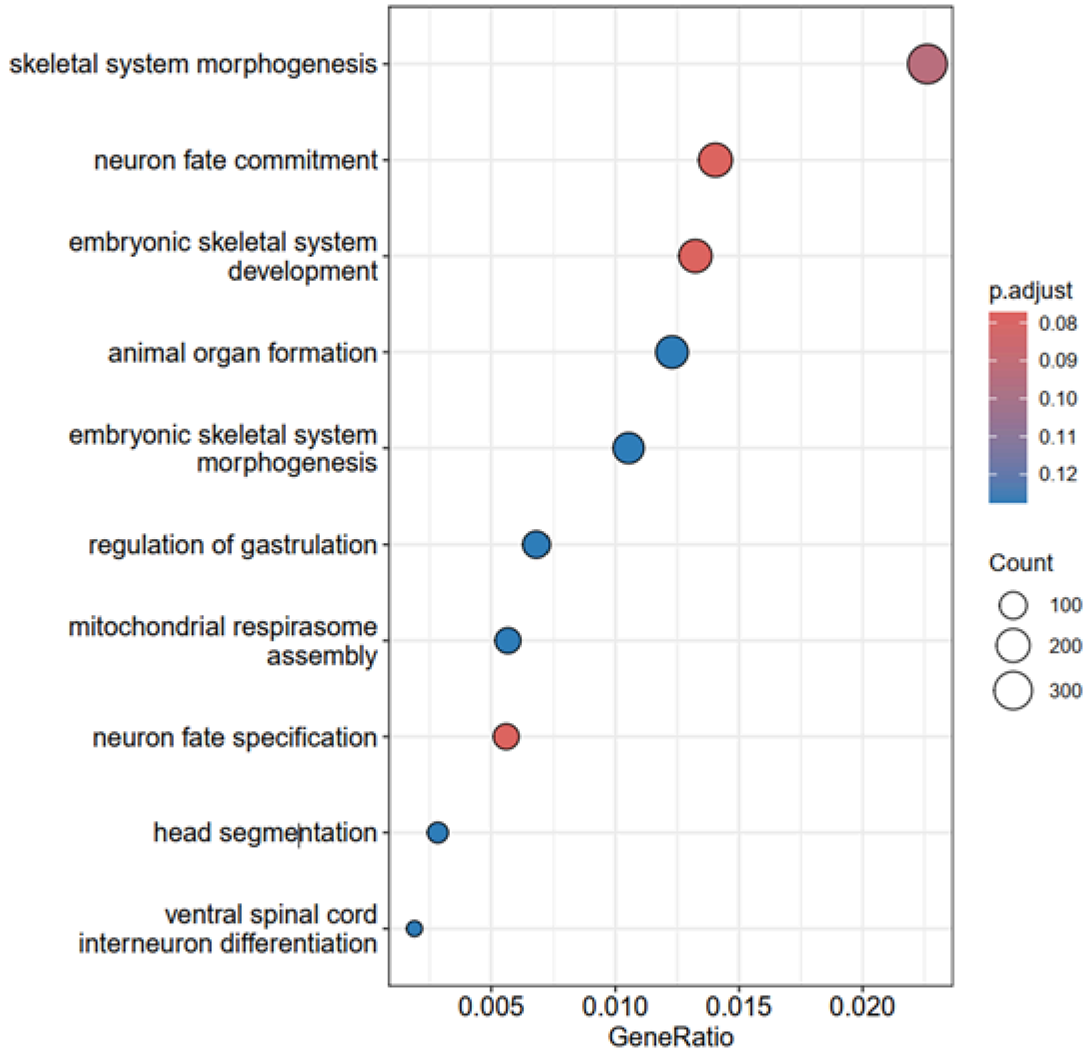

**Supplementary Fig. 2.** The GO enrichment of softcore genes

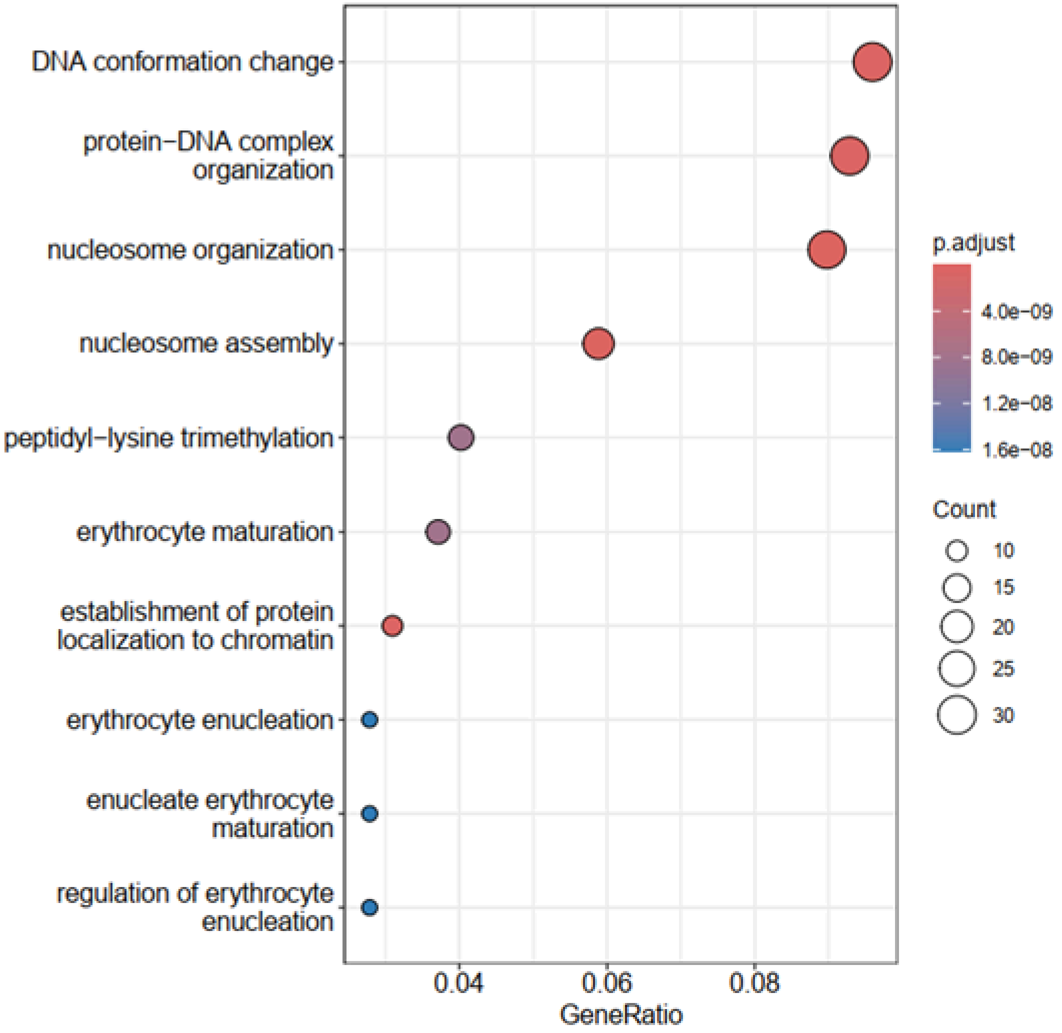

**Supplementary Fig. 3.** The GO enrichment of dispensable genes

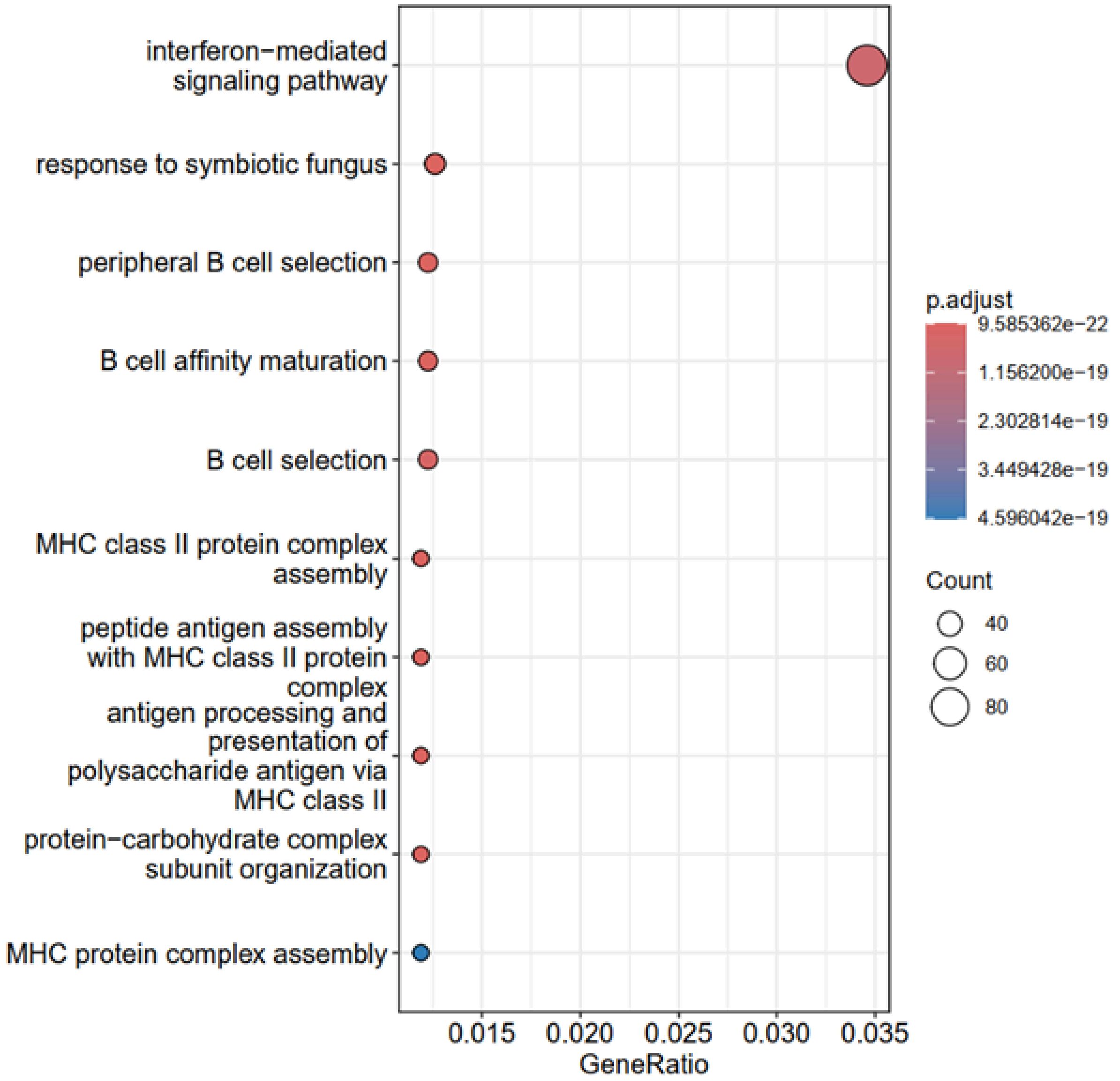

**Supplementary Fig. 4.** The GO enrichment of Private genes

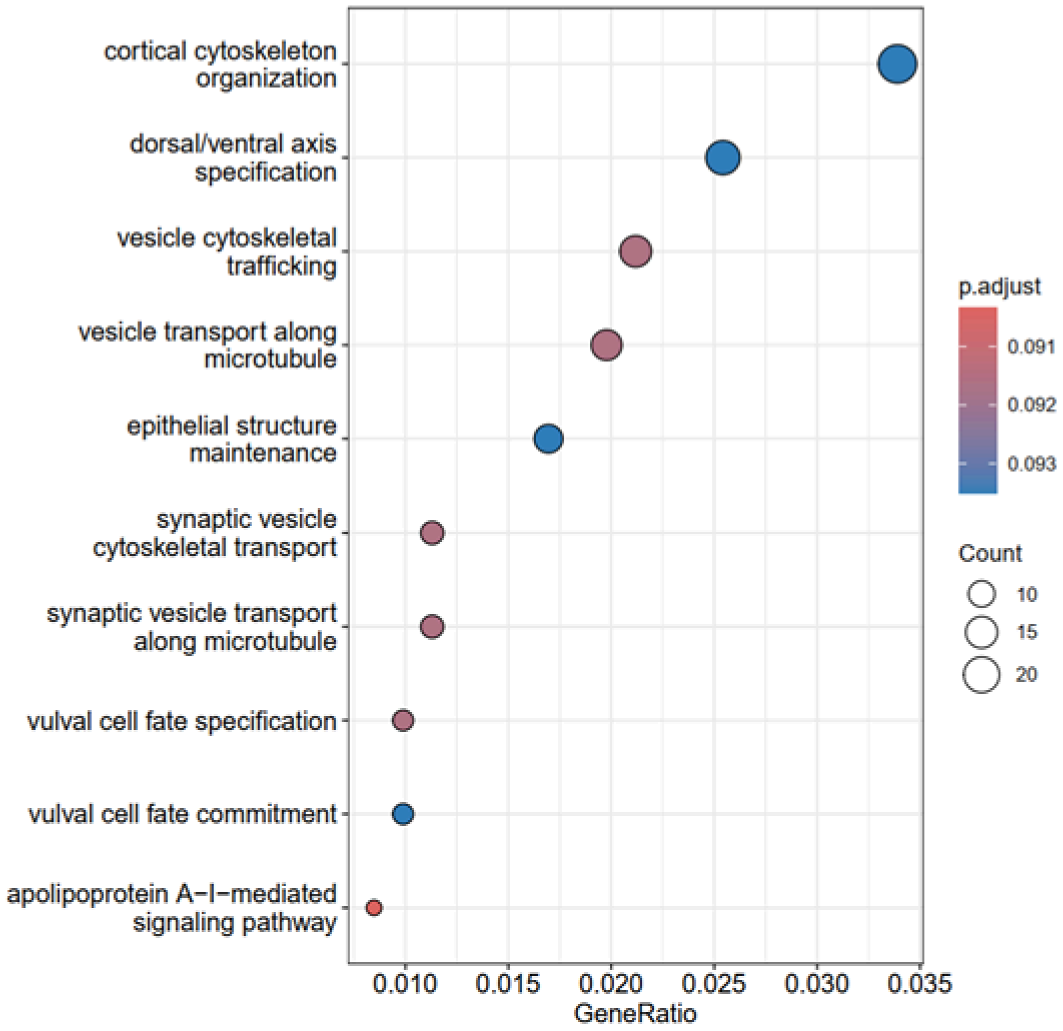

**Supplementary Fig. 5.** The gene count in 45 nine-spined stickleback populations

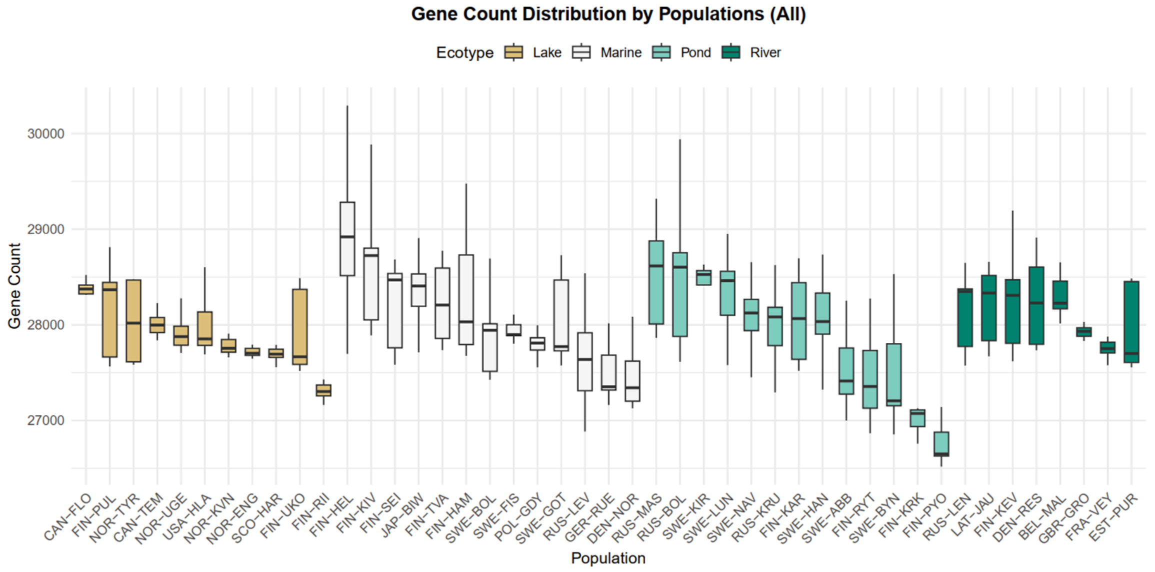

**Supplementary Fig. 6.** The mean number of genes lost in 40 nine-spined stickleback populations. Bars are colored according to ecotype: blue represents the marine ecotype, green represents the pond ecotype, pink represents the river ecotype, and yellow represents the lake ecotype.

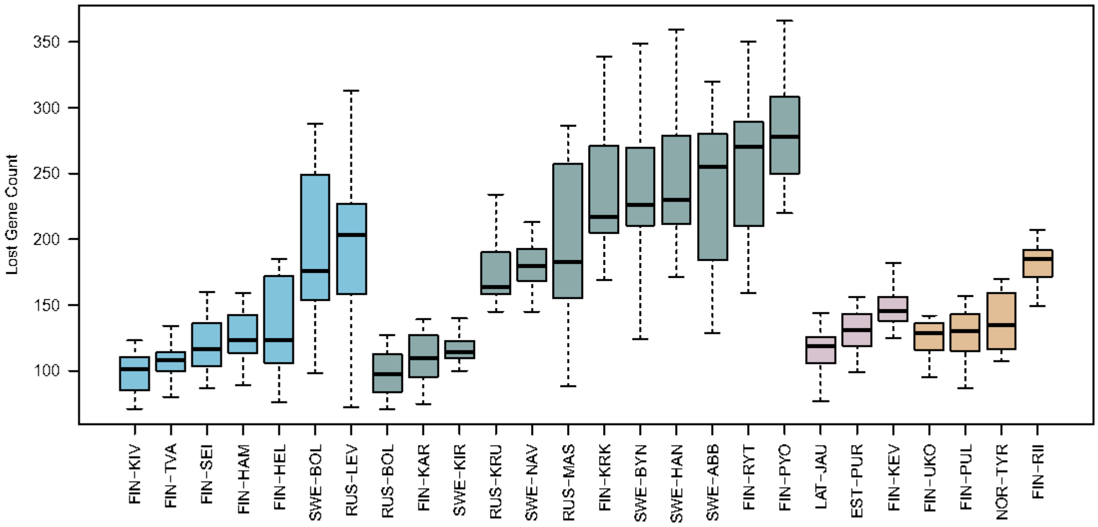

**Supplementary Fig. 7.** Comparison neutrality index and direction of selection between core and PAV genes.

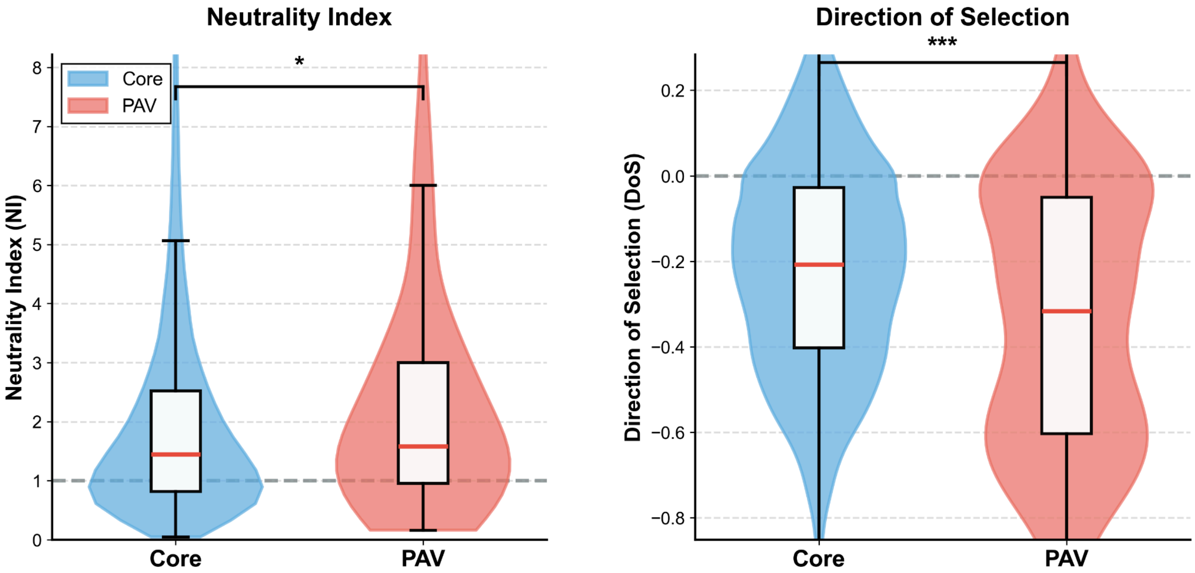

**Supplementary Fig. 8** Freshwater-biased gene loss and descriptive gene-level differentiation summaries.

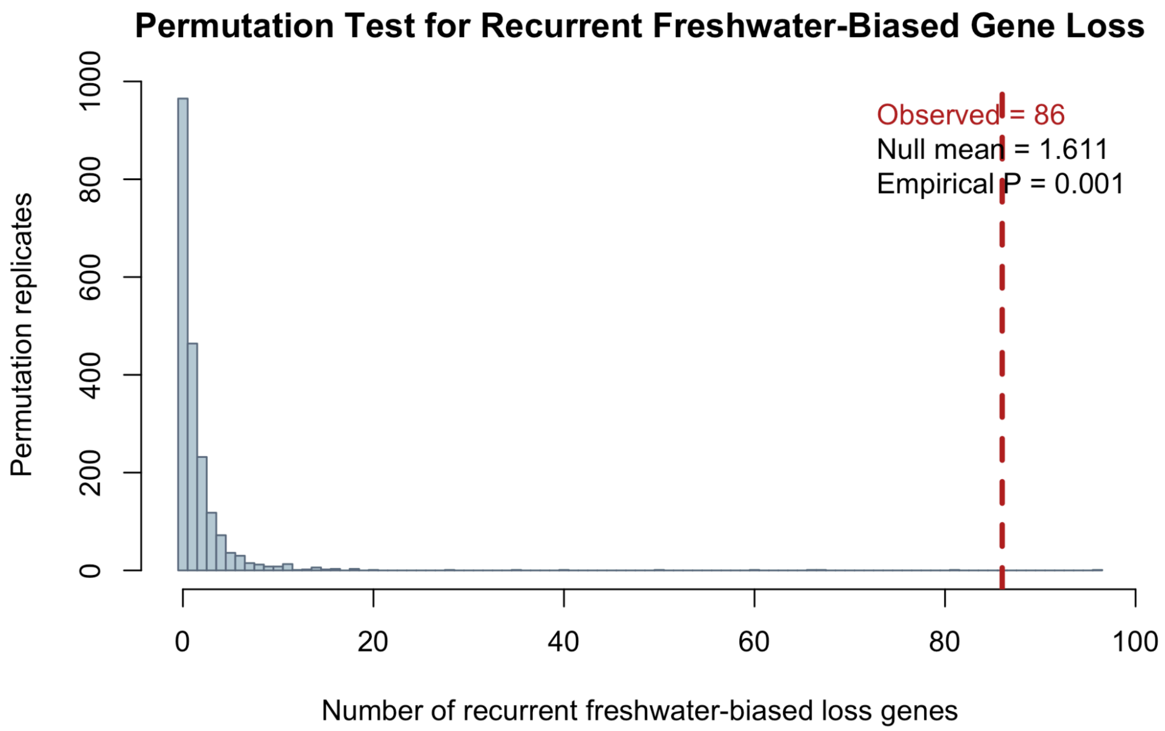

Permutation Test for Recurrent Freshwater-Biased Gene Loss
Observed = 86
Null mean = 1.611
Empirical P = 0.001
Permutation replicates
0
200
400
600
800
1000
0
20
40
60
80
100
Number of recurrent freshwater-biased loss genes

**Supplementary Fig. 9** Recurrent gene loss in freshwater environments remains consistent across multiple populations, even when applying increasingly strict criteria for parallel evolution.

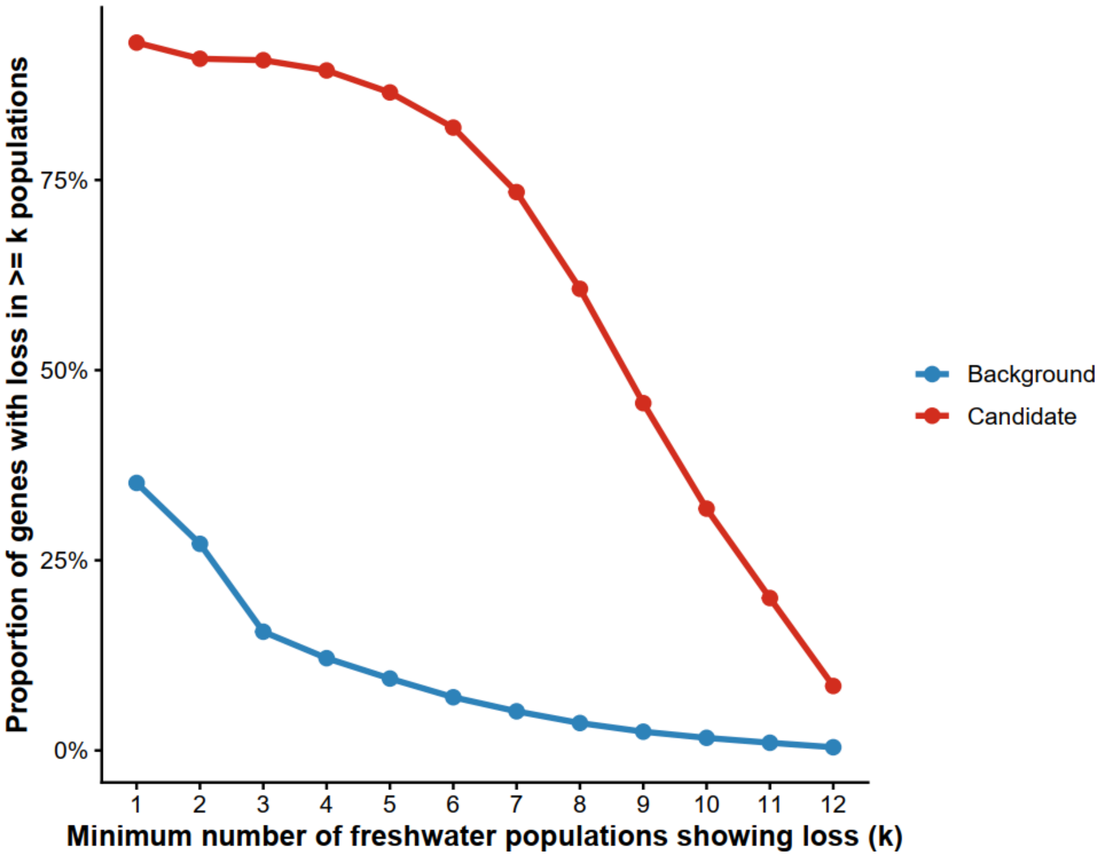

**Supplementary Fig. 10** Genes exhibiting significant, freshwater-associated loss across all 12 populations of nine-spined sticklebacks. Dots denote significant gene loss events specific to individual populations.

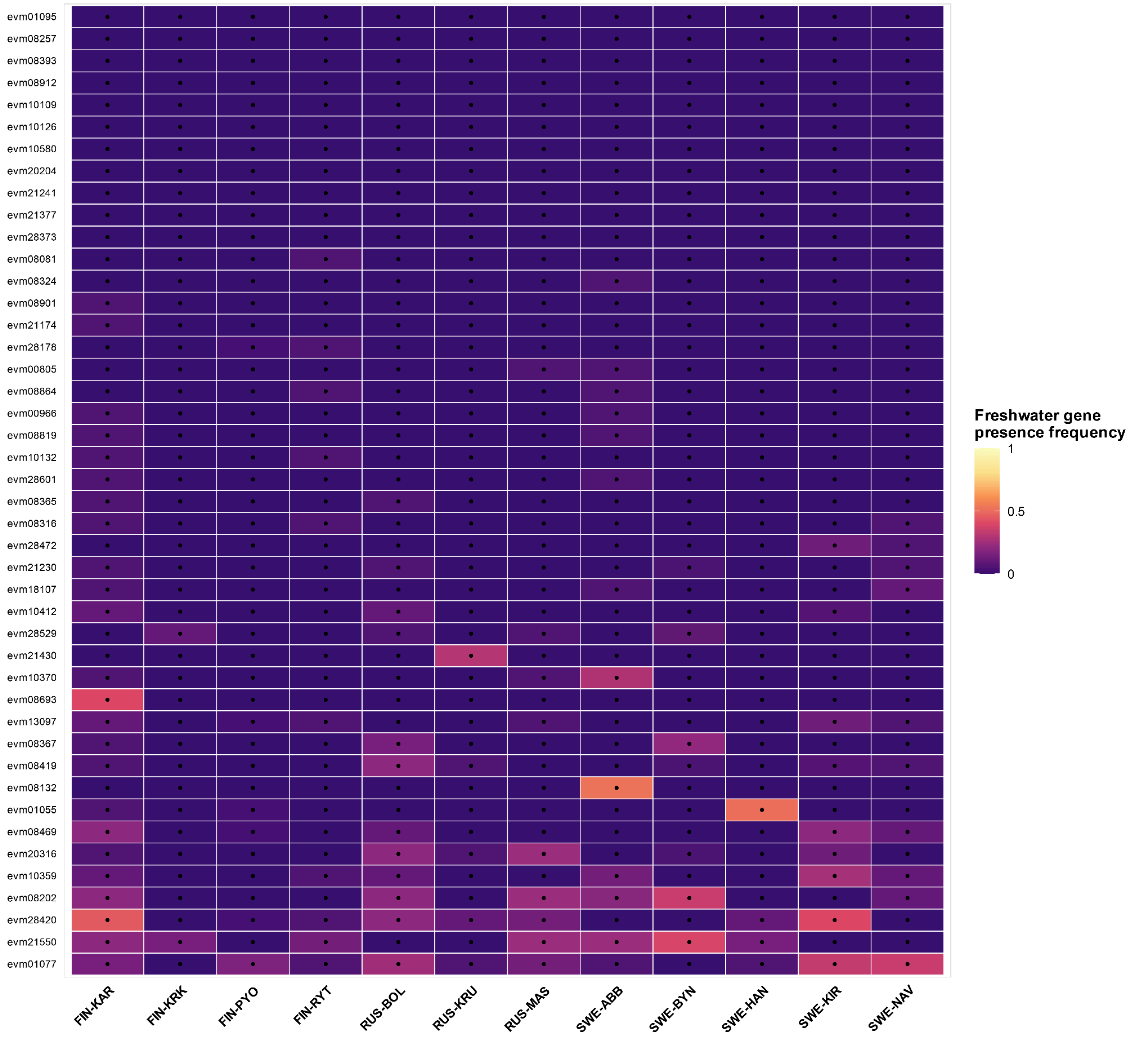

**Supplementary Table 1a. Sample sets used in each analysis.**

| Analysis | Sample set | n | Populations | Sequencing | Purpose |
|---|---|---|---|---|---|
| Pangenome construction | All re-sequenced genomes | 1,478 | 47 | 5-80x | *De novo* assembly and map-to-pan non-reference sequence recovery |
| Pangenome/PAV modeling | 887 wild samples + 80 F0 high-depth samples | 967 | 45 wild + 8 high-depth populations | ~10x and ~80x | Pangenome and core-genome growth curves |
| Population-level PAV and SNP analyses | 887 wild unrelated individuals | 887 | 45 | ~10x | PAV frequencies, SNP calling, LoF/MK summaries, descriptive FST |
| Ancestral-gene-loss subset | 809 wild unrelated non-outgroup individuals | 809 | 40 | ~10x | Main gene-loss models and sensitivity analyses |
| Assembly-size analysis | 434 high-depth individuals | 434 | 8 (4 pond, 4 marine) | ~80x | Short-read assembly size, repeat content, and LMMs |

Note: the main ancestral-gene-loss models used all 40 non-outgroup populations. The DepthMean/DepthMedian coverage robustness models used the 39 populations with available whole-gene depth summaries (SWE-LUN excluded); the text-based excov coverage models retained all 40 populations.

**Supplementary Table 2. Information on gene annotations in different genome assemblies and Non-reference-represented (NRR) assemblies identified in this study.**

| Accesstion | BUSCO（%） | Contig number | Gene number |
|---|---|---|---|
| v8 raw annotation | 92.5 | 129 | 26803 |
| v8 update annotation | 96.1 | 129 | 27450 |
| NRRs annotation | 0.7 | 263128 | 11151 |

**Supplementary Table 3. The number and proportion of genes in different categories.**

| Class | Frequency | Proportion |
|---|---|---|
| Core | 24130 | 0.667 |
| Softcore | 2009 | 0.056 |
| Dispensable | 9044 | 0.250 |
| Private | 991 | 0.027 |

**Supplementary Table 4. Chromosome-specific distribution of gene categories.**

| Chromosome | Type | Core (%) | Softcore (%) | Dispensable (%) |
|---|---|---|---|---|
| LG1-LG11, LG13-LG21 | Auto | 89.9–95.3 | 3.9–10.0 | 0.8–2.9 |

| | | | | |
|---|---|---|---|---|
| LG12 | X | 86.8 | 10.1 | 3.1 |
| LG22 | Y | 5.1 | 18.5 | 76.4 |

**Supplementary Table 5. Number of genes in different pangenome gene categories in the reference assembly and NRRs**

| **Accesstion** | **Core** | **Softcore** | **Dispensable** | **Private** |
|---|---|---|---|---|
| Reference | 24121 | 1931 | 1396 | 0 |
| NRRs | 9 | 78 | 7648 | 991 |

**Supplementary Table 6. Lineage comparison and fixed differences in PAV frequencies between the EL and WL lineage**

| Metric | Value | Interpretation |
|---|---|---|
| Total PAV genes compared between EL and WL | 12,044 | Full lineage-level comparison set |
| Fully fixed diagnostic PAVs between EL and WL | 0 | No absolute lineage-specific presence/absence markers were recovered |

**Supplementary Table 7 Gene-loss robustness and sensitivity analyses**

| Analysis | Model / response | Population-size term | Ecotype term | Additional | Interpretation |
|---|---|---|---|---|---|

| | | | | | |
|---|---|---|---|---|---|
| Main model | Phylogenetic Poisson mixed model | beta = -0.0968, 95% CI = -0.1284 to -0.0655, pMCMC < 0.00012 | Freshwater vs marine beta = -0.2864, 95% CI = -0.3420 to -0.2285, pMCMC < 0.00012 | Phylogeny | Gene loss declines with larger Ne |
| Lineage sensitivity | Main model plus lineage term | Negative, similar to main model | Similar to main model | Lineage: weak, CI overlaps 0 | Lineage does not overturn demographic effect |
| Population-level Gaussian | MeanLostGenes ~ log10(Ne + 1) + EcotypeSimple | beta = -100.95, P = 3.89 × 10^-4 | Marine beta = 14.86, P = 0.369 | None | Mean gene loss tracks Ne, not ecotype |
| Population-level Gaussian | MedianLostGenes ~ log10(Ne + 1) + EcotypeSimple | beta = -96.94, P = 0.00106 | Marine beta = 14.72, P = 0.399 | None | Median summary agrees |
| Coverage robustness | MeanLostGenes ~ log10(Ne + 1) + EcotypeSimple + DepthMean | beta = -60.84, P = 0.00812 | Marine P = 0.410 | Depth: beta = -0.0673, P = 1.16 × 10^-5 | Ne effect survives depth control |
| Coverage robustness | MedianLostGenes ~ log10(Ne + 1) + EcotypeSimple + DepthMean | beta = -52.69, P = 0.0220 | Marine P = 0.435 | Depth: beta = -0.0745, P = 2.80 × 10^-6 | Median result robust to depth |
| Coverage robustness | MeanLostTxt ~ log10(Ne + 1) + EcotypeSimple + MeanGeneDepthAllGenes | beta = -119.70, P = 4.60 × 10^-8 | Marine P = 0.102 | Gene depth: beta = -15.32, P = 4.03 × 10^-8 | Text-based losses agree |
| Coverage robustness | MeanLostTxt ~ log10(Ne + 1) + EcotypeSimple + MeanNotCoveredFraction | beta = -83.65, P = 5.48 × 10^-4 | Marine P = 0.679 | Not-covered: beta = 4310.63, P = 2.21 × 10^-4 | Ne association persists |
| Alternative Ne estimator | MeanLostGenes ~ log10(MSMC2 Ne + 1) + EcotypeSimple | beta = -61.20, P = 2.68 × 10^-4 | Marine beta = 19.68, P = 0.247 | None | Not specific to Ne estimator |
| Weighted count model | Negative binomial TotalLost ~ log10(MSMC2 Ne + 1) + EcotypeSimple + offset(log N) | beta = -0.3527, P = 6.12 × 10^-5 | Marine beta = 0.120, P = 0.215 | Offset log N | Count model supports demographic effect |

| | | | | | |
|---|---|---|---|---|---|
| Weighted count model | Negative binomial TotalLost ~ log10(Ne + 1) + EcotypeSimple + offset(log N) | beta = -0.5801, P = 1.06 × 10^-4 | Marine beta = 0.0899, P = 0.342 | Offset log N | Same under Ne count model |
| Rarefaction sensitivity | Common rarefied sample size n = 8 | Ne beta median = -100.73, 95% interval = [-110.86, -90.88] | Marine median = 14.65, 95% interval = [8.16, 21.81] | Rarefied n = 8 | Ne effect stable after rarefaction |
| Fixed-loss partition | Negative binomial FixedLoss ~ log10(Ne + 1) + EcotypeSimple | beta = -0.0970, P = 0.184 | Marine beta = -0.0154, P = 0.739 | None | Fixed loss does not drive signal |
| Segregating-loss partition | Negative binomial SegregatingLoss ~ log10(Ne + 1) + EcotypeSimple | beta = 0.2740, P = 0.0766 | Marine beta = 0.0912, P = 0.350 | None | Segregating loss heterogeneous; no ecotype effect |
| Locality heterogeneity | Additive vs interaction model for Lost_gene_count | Not primary term | Interaction EcotypeSimple x Locality: P = 7.647 × 10^-5 | 3 localities | Geographically heterogeneous |

### Supplementary Table 8 Exon-level coverage support for recurrent freshwater-loss candidates

Group-level summary:

| Group | Records | Median fraction of exons covered | Median exon depth | Proportion with zero exon depth | Dominant coverage category |
|---|---|---|---|---|---|
| Freshwater_LOST | 1,521 | 0.000 | 0.000 | 0.971 | not_covered |
| Freshwater_PRESENT | 643 | 1.000 | 7.031 | 0.000 | 0-10 |
| Marine_LOST | 383 | 0.000 | 0.000 | 0.932 | not_covered |
| Marine_PRESENT | 693 | 1.000 | 6.189 | 0.000 | 0-10 |

Gene-level summary for the four traced candidates with direct freshwater-loss versus marine-presence contrasts:

| GeneID | Group | Records | Populations | Median fraction of exons covered | Median exon depth | Proportion with zero exon depth | Dominant coverage category |
|---|---|---|---|---|---|---|---|
| evm00966 | Freshwater_LOST | 387 | 23 | 0.000 | 0.000 | 0.956 | not_covered |

| | | | | | | | |
|---|---|---|---|---|---|---|---|
| evm00966 | Freshwater_PRESENT | 154 | 15 | 0.978 | 4.741 | 0.000 | 0-10 |
| evm00966 | Marine_LOST | 120 | 12 | 0.000 | 0.000 | 0.933 | not_covered |
| evm00966 | Marine_PRESENT | 149 | 11 | 0.940 | 4.151 | 0.000 | 0-10 |
| evm08367 | Freshwater_LOST | 336 | 22 | 0.000 | 0.000 | 0.970 | not_covered |
| evm08367 | Freshwater_PRESENT | 205 | 17 | 0.973 | 6.792 | 0.000 | 0-10 |
| evm08367 | Marine_LOST | 67 | 9 | 0.000 | 0.000 | 0.791 | not_covered |
| evm08367 | Marine_PRESENT | 202 | 12 | 0.914 | 7.324 | 0.000 | 0-10 |
| evm08819 | Freshwater_LOST | 343 | 20 | 0.000 | 0.000 | 0.956 | not_covered |
| evm08819 | Freshwater_PRESENT | 198 | 14 | 1.000 | 10.843 | 0.000 | 10-20 |
| evm08819 | Marine_LOST | 81 | 10 | 0.000 | 0.000 | 0.975 | not_covered |
| evm08819 | Marine_PRESENT | 188 | 11 | 1.000 | 7.837 | 0.000 | 0-10 |
| evm08864 | Freshwater_LOST | 455 | 27 | 0.000 | 0.000 | 0.996 | not_covered |
| evm08864 | Freshwater_PRESENT | 86 | 10 | 0.972 | 4.099 | 0.000 | 0-10 |
| evm08864 | Marine_LOST | 115 | 12 | 0.000 | 0.000 | 0.983 | not_covered |
| evm08864 | Marine_PRESENT | 154 | 11 | 1.000 | 5.585 | 0.000 | 0-10 |

### Supplementary Table 9. Comparison of pangenome studies across animal species.

| Species | Taxonomic | Sample Size | Number of Populations / Lineages | Novel sequence Mb | Novel sequence % | Reference |
|---|---|---|---|---|---|---|
| *Pungitius pungitius* | Fish | 1,478 | 47 populations | 234.15 Mb | 44% | This study |
| *Bos grunniens* (Yak) | Mammal | 366 (16 *de novo* + 350 resequenced) | 51 breeds (Wild and Domestic) | 290 Mb | - | Lan et al., 2024 |
| *Sus scrofa* (Pig) | Mammal | 250 | 32 breeds | 308.3 Mb | - | Li et al., 2023 |
| *Aphelocoma* spp. (Scrub Jays) | Bird | 45 individuals (90 haplotypes) | 3 species | 7,300 Mb (7.3 Gb) | ~6.5% | Edwards et al., 2025 |
| *Haemorhous mexicanus* (House Finch) | Bird | 16 individuals (32 haplotypes) | 3 lineages | - | - | Fang & Edwards, 2024 |
| *Apis cerana* (Asian Honeybee) | Insect | 526 (1 ref + 525 resequenced) | 9 populations | 127.6 Mb | ~58.6% | Li et al., 2024 |

| | | | | | | |
|---|---|---|---|---|---|---|
| *Lake Malawi cichlids* (Haplochromine cichlids) | Fish | 8 assemblies | 7 species | 291.47 Mb | 33.1% | Quah et al., 2025 |
| *Bos taurus* (Cattle) | Mammal | 898 | 57 breeds | 83 Mb | 3.1% | Zhou et al., 2022 |
| *Ovis aries* (Sheep) | Mammal | 15 | 13 breeds | 130.3 Mb | - | Li et al., 2023 |

Supplementary Table 10 Broader recurrent freshwater-loss candidate genes from the descriptive follow-up screen

| GeneID | SeqID | Start | End | Strand | Gene Length | FW absent | FW total | FW absent rate | MAR absent | MAR total | MAR absent rate | FW-MAR diff | FW pops any loss | MAR pops any loss | FW pops >= 50% absent | MAR pops >= 50% absent | GC content | Repeat overlap fraction |
|---|---|---|---|---|---|---|---|---|---|---|---|---|---|---|---|---|---|---|
| evm13097 | fin_por11_F1A04_contig18168 | 18 | 182 | - | 165 | 347 | 540 | 0.642593 | 70 | 269 | 0.260223 | 0.38237 | 25 | 12 | 16 | 1 | 0.5394 | 0.0 |
| evm28529 | umeinb12_F2_022_contig54440 | 69 | 949 | + | 881 | 442 | 540 | 0.818519 | 143 | 269 | 0.531599 | 0.28692 | 27 | 12 | 24 | 4 | 0.4506 | 0.0 |
| evm28373 | swe_ume22_F0_M_contig172892 | 135 | 550 | + | 416 | 432 | 540 | 0.8 | 104 | 269 | 0.386617 | 0.413383 | 28 | 12 | 22 | 3 | 0.5024 | 0.1538 |
| evm10580 | SWE-LUN-40.scaffold133975_1 | 47 | 464 | + | 418 | 383 | 540 | 0.709259 | 152 | 269 | 0.565056 | 0.144203 | 23 | 12 | 19 | 8 | 0.4545 | 0.0 |
| evm21230 | fin_tva32_F1B03_contig61663 | 1 | 597 | - | 597 | 427 | 540 | 0.790741 | 132 | 269 | 0.490706 | 0.300034 | 26 | 12 | 22 | 5 | 0.5126 | 0.0 |
| evm21430 | fin_tva36_F1B07_contig162632 | 979 | 1711 | - | 733 | 382 | 540 | 0.707407 | 69 | 269 | 0.256506 | 0.450902 | 25 | 11 | 20 | 1 | 0.513 | 0.0 |
| evm08393 | GBR-GRO-16.scaffold89404_2 | 200 | 356 | - | 157 | 366 | 540 | 0.677778 | 106 | 269 | 0.394052 | 0.283726 | 23 | 11 | 17 | 2 | 0.5414 | 0.0 |

| | | | | | | | | | | | | | | | | | | |
|---|---|---|---|---|---|---|---|---|---|---|---|---|---|---|---|---|---|---|
| evm 28178 | swe_ume05_F0_M_contig105346 | 67 | 685 | + | 619 | 410 | 540 | 0.759259 | 159 | 269 | 0.591078 | 0.168181 | 24 | 12 | 21 | 9 | 0.588 | 0.0275 |
| evm 00966 | DEN-RES-17.C10192842_1 | 87 | 270 | - | 184 | 398 | 540 | 0.737037 | 132 | 269 | 0.490706 | 0.246331 | 25 | 12 | 19 | 4 | 0.5272 | 0.0 |
| evm 21377 | fin_tva36_F0_M_contig125332 | 218 | 535 | + | 318 | 432 | 540 | 0.8 | 166 | 269 | 0.6171 | 0.1829 | 25 | 12 | 23 | 9 | 0.5252 | 0.0 |
| evm 21550 | porout03_F2_05_contig9625 | 13 | 170 | - | 158 | 381 | 540 | 0.705556 | 74 | 269 | 0.275093 | 0.430463 | 27 | 10 | 22 | 3 | 0.5696 | 0.0 |
| evm 00805 | CAN-TEM-24.scaffold1266_1 | 102 | 528 | + | 427 | 473 | 540 | 0.875926 | 154 | 269 | 0.572491 | 0.303435 | 28 | 12 | 24 | 7 | 0.466 | 0.0 |
| evm 08367 | GBR-GRO-12.C1123469_1 | 63 | 358 | - | 296 | 369 | 540 | 0.683333 | 108 | 269 | 0.401487 | 0.281846 | 25 | 11 | 17 | 5 | 0.4561 | 0.0 |
| evm 20316 | fin_raa13_F1B062_contig39682 | 167 | 374 | - | 208 | 437 | 540 | 0.809259 | 108 | 269 | 0.401487 | 0.407772 | 27 | 12 | 24 | 1 | 0.5865 | 0.0 |
| evm 08419 | GBR-GRO-3.scaffold74362_1 | 98 | 447 | + | 350 | 305 | 540 | 0.564815 | 75 | 269 | 0.27881 | 0.286004 | 22 | 11 | 15 | 1 | 0.5371 | 0.1229 |
| evm 10370 | SWE-GOT-16.scaffold123605_1 | 389 | 771 | - | 383 | 391 | 540 | 0.724074 | 93 | 269 | 0.345725 | 0.378349 | 26 | 12 | 19 | 1 | 0.5222 | 0.0 |
| evm 01055 | FIN-HEL-19-f.scaffold51857_1 | 165 | 425 | - | 261 | 345 | 540 | 0.638889 | 55 | 269 | 0.204461 | 0.434428 | 21 | 10 | 17 | 1 | 0.567 | 0.0 |
| evm 08132 | FIN-TVA-17.scaffold15721_2 | 84 | 290 | - | 207 | 329 | 540 | 0.609259 | 37 | 269 | 0.137546 | 0.471713 | 20 | 6 | 15 | 1 | 0.4686 | 0.0 |
| evm 08257 | FRA-VEY-11.scaffold7654_1 | 225 | 2885 | - | 2661 | 430 | 540 | 0.796296 | 103 | 269 | 0.3829 | 0.413397 | 26 | 11 | 22 | 3 | 0.4476 | 0.46116 |
| evm 10126 | SCO-HAR-9.scaffold3081_1 | 139 | 379 | + | 241 | 373 | 540 | 0.690741 | 46 | 269 | 0.171004 | 0.519737 | 21 | 6 | 18 | 1 | 0.3734 | 0.0 |
| evm 10132 | SCO-HAR-9.scaffold99744_1 | 1330 | 1698 | - | 369 | 446 | 540 | 0.825926 | 102 | 269 | 0.379182 | 0.446744 | 28 | 12 | 23 | 2 | 0.439 | 0.0 |
| evm 08819 | NOR-UGE-26.C2098035_1 | 131 | 459 | + | 329 | 345 | 540 | 0.638889 | 86 | 269 | 0.319703 | 0.319186 | 21 | 10 | 16 | 2 | 0.5076 | 0.08216 |

| | | | | | | | | | | | | | | | | | | |
|---|---|---|---|---|---|---|---|---|---|---|---|---|---|---|---|---|---|---|
| evm10109 | SCO-HAR-7.scaffold1052_1 | 299 | 196 | - | 168 | 444 | 540 | 0.822222 | 111 | 269 | 0.412639 | 0.409583 | 23 | 10 | 23 | 4 | 0.5357 | 0.0 |
| evm08693 | NOR-ENG-2.scaffold109179_1 | 288 | 507 | - | 220 | 353 | 540 | 0.653704 | 63 | 269 | 0.234201 | 0.419503 | 21 | 9 | 17 | 1 | 0.5455 | 0.0 |
| evm08901 | POL-GDY-26.scaffold158851_2 | 161 | 540 | - | 380 | 385 | 540 | 0.712963 | 57 | 269 | 0.211896 | 0.501067 | 24 | 11 | 19 | 1 | 0.55 | 0.0 |
| evm21241 | fin_tva32_F1B04_contig163331 | 144 | 556 | + | 413 | 435 | 540 | 0.805556 | 82 | 269 | 0.304833 | 0.500723 | 23 | 9 | 22 | 3 | 0.4794 | 0.0 |
| evm01077 | FIN-HEL-36-f.scaffold2078_1 | 173 | 333 | - | 161 | 362 | 540 | 0.67037 | 57 | 269 | 0.211896 | 0.458474 | 23 | 6 | 22 | 3 | 0.4224 | 0.0 |
| evm08912 | POL-GDY-7.scaffold175572_1 | 89 | 569 | - | 481 | 372 | 540 | 0.688889 | 75 | 269 | 0.27881 | 0.410078 | 21 | 9 | 18 | 1 | 0.5156 | 0.0 |
| evm20204 | fin_raa04_F1B03_contig43200 | 456 | 686 | - | 231 | 344 | 540 | 0.637037 | 49 | 269 | 0.182156 | 0.454881 | 18 | 5 | 16 | 1 | 0.5671 | 0.0 |
| evm08324 | FRA-VEY-3.scaffold3660_1 | 195 | 518 | - | 324 | 369 | 540 | 0.683333 | 54 | 269 | 0.200743 | 0.48259 | 22 | 10 | 17 | 1 | 0.4074 | 0.0 |
| evm28472 | umeinb11_F2_01_contig86900 | 177 | 368 | - | 192 | 444 | 540 | 0.822222 | 110 | 269 | 0.408922 | 0.4133 | 24 | 11 | 24 | 4 | 0.4792 | 0.0 |
| evm08202 | FIN-UKO-2.scaffold31792_1 | 72 | 743 | + | 672 | 324 | 540 | 0.6 | 88 | 269 | 0.327138 | 0.272862 | 24 | 12 | 17 | 3 | 0.5432 | 0.0 |
| evm08081 | FIN-SEI-1.scaffold142733_2 | 703 | 1082 | - | 380 | 383 | 540 | 0.709259 | 53 | 269 | 0.197026 | 0.512233 | 24 | 11 | 19 | 1 | 0.55 | 0.0 |
| evm08365 | GBR-GRO-11.scaffold42547_1 | 40 | 572 | + | 533 | 407 | 540 | 0.753704 | 57 | 269 | 0.211896 | 0.541808 | 27 | 11 | 20 | 1 | 0.4878 | 0.0 |
| evm08864 | POL-GDY-11.C1554172_1 | 205 | 365 | - | 161 | 469 | 540 | 0.868519 | 132 | 269 | 0.490706 | 0.377812 | 27 | 12 | 25 | 6 | 0.4099 | 0.0 |
| evm28420 | swe_ume34_F0_M_contig72965 | 46 | 216 | - | 171 | 288 | 540 | 0.533333 | 51 | 269 | 0.189591 | 0.343742 | 19 | 10 | 15 | 1 | 0.6082 | 0.0 |
| evm08469 | GER-RUE-19.scaffold46992_1 | 289 | 456 | - | 168 | 342 | 540 | 0.633333 | 87 | 269 | 0.32342 | 0.309913 | 24 | 11 | 17 | 1 | 0.5238 | 0.0 |

| | | | | | | | | | | | | | | | | | | |
|---|---|---|---|---|---|---|---|---|---|---|---|---|---|---|---|---|---|---|
| evm10359 | SWE-GOT-10.scaffold148333_1 | 248 | 436 | + | 189 | 335 | 540 | 0.62037 | 63 | 269 | 0.234201 | 0.38617 | 22 | 11 | 16 | 1 | 0.6296 | 0.0 |
| evm21174 | fin_tva32_F1A02_contig66117 | 922 | 1071 | - | 150 | 414 | 540 | 0.766667 | 144 | 269 | 0.535316 | 0.231351 | 27 | 12 | 21 | 6 | 0.5133 | 0.0 |
| evm10412 | SWE-GOT-7.scaffold155282_1 | 161 | 346 | + | 186 | 323 | 540 | 0.598148 | 60 | 269 | 0.223048 | 0.3751 | 23 | 11 | 16 | 1 | 0.6022 | 0.0 |
| evm08316 | FRA-VEY-2.scaffold54721_1 | 46 | 780 | + | 735 | 368 | 540 | 0.681481 | 166 | 269 | 0.6171 | 0.064381 | 23 | 11 | 19 | 9 | 0.4925 | 0.3293 |
| evm01095 | FIN-HEL-78-m-2.scaffold672_1 | 319 | 486 | - | 168 | 437 | 540 | 0.809259 | 88 | 269 | 0.327138 | 0.482122 | 25 | 11 | 22 | 3 | 0.5238 | 0.0 |
| evm28601 | umeout09_F2_09_contig148187 | 334 | 586 | - | 253 | 350 | 540 | 0.648148 | 61 | 269 | 0.226766 | 0.421382 | 24 | 9 | 17 | 1 | 0.5336 | 0.0 |
| evm18107 | fin_por12_F0_M_contig2237 | 60 | 362 | + | 303 | 420 | 540 | 0.777778 | 131 | 269 | 0.486989 | 0.290789 | 27 | 12 | 22 | 5 | 0.4521 | 0.0 |
